# Ancestral P-body proteins rewired for autophagic recycling in the early land plant *Marchantia polymorpha*

**DOI:** 10.1101/2025.08.09.669463

**Authors:** Alibek Abdrakhmanov, Aleksandra S. Anisimova, Ranjith K. Papareddy, Nenad Grujic, Elizabeth Ethier, Marion Clavel, G. Elif Karagöz, Erinc Hallacli, Yasin Dagdas

## Abstract

Processing bodies (P-bodies) are conserved ribonucleoprotein (RNP) granules central to RNA metabolism across eukaryotes. Although the mechanisms underlying their assembly are well understood, the pathways governing their selective turnover remain unclear. Here, we identify the conserved decapping proteins EDC4 and DCP1 as a selective autophagy receptor pair responsible for P-body turnover in the early land plant *Marchantia polymorpha*. MpEDC4 engages ATG8 via a canonical AIM motif, while MpDCP1 contains a previously unrecognized reverse AIM within its intrinsically disordered region. Mutations disrupting these motifs impair autophagic degradation of P-bodies, demonstrating a cooperative receptor mechanism. Notably, this autophagic function is lineage-specific, as orthologs in *Arabidopsis* and humans lack ATG8-binding capacity. Strikingly, heterologous expression of MpEDC4 in human cells promotes degradation of α-synuclein, a protein strongly linked to Parkinson’s disease etiology. Our findings thus uncover an evolutionary innovation that links RNA metabolism to selective autophagy and opens avenues for cross-kingdom engineering of targeted protein degradation pathways.

## Introduction

Processing bodies (P-bodies) are membraneless cytoplasmic ribonucleoprotein (RNP) granules that regulate mRNA metabolism by controlling RNA decay, translational repression, and sequestration^1,2^. These dynamic structures consist of enzymes, such as decapping and deadenylation complexes, RNA-binding proteins, and translational repressors that maintain mRNA homeostasis as well as enable rapid modulation of gene expression in response to environmental stress and developmental signals^3^. P-body assembly and disassembly are governed by multivalent interactions between proteins and RNAs, bridged by intrinsically disordered regions (IDRs) of protein components^4,5^. The formation of RNP granules requires both non-translated RNAs and RNA-binding proteins, many of which contain aggregation-prone prion-like domains or low-complexity intrinsically disordered regions^4,5^. Mutations in these proteins are frequently associated with neurodegenerative diseases such as Alzheimer’s disease, Parkinson’s disease, frontotemporal dementia, amyotrophic lateral sclerosis, and spinal muscular atrophy^6^. Therefore, precise control of P-body dynamics is crucial for maintaining cellular homeostasis. Recent evidence suggests that the regulated turnover of RNPs occurs not only through controlled assembly and disassembly but also via targeted degradation pathways^7–9^. Among these, selective autophagy is emerging as a potential mechanism that may mediate the specific recycling of RNP granules.

Macroautophagy (hereafter autophagy) is a conserved catabolic process responsible for the degradation and recycling of cellular components. Double-membrane vesicles called autophagosomes encapsulate cytoplasmic cargo and deliver it to lysosomes (in animals) or vacuoles (in plants or yeast) for degradation^10–12^. Autophagosome formation and maturation are orchestrated by evolutionarily conserved Autophagy-related (ATG) proteins^13^. Selective autophagy is a specialized form of this process, whereby specific cargoes are recognized and targeted for degradation by selective autophagy receptors (SARs)^14^. SARs act by linking autophagic cargo to the autophagy machinery through direct interactions with the core autophagy protein ATG8 via ATG8-interacting motifs (AIMs)^15^. AIMs are short linear motifs, typically following the consensus sequence [F/W/Y]xx[I/L/V], where x represents any amino acid^16^. Diverse cellular substrates are selectively degraded through autophagy, including damaged organelles^17–20^, protein aggregates^21,22^, pathogens^23,24^, and membraneless organelles like RNP granules^25^ or ribosomes^26,27^. However, despite extensive knowledge of selective autophagy mechanisms, the receptors mediating P-body recycling remain unknown. Here, we investigated the molecular mechanisms underlying selective autophagic recycling of P-bodies using the liverwort *Marchantia polymorpha* as a model system.

## Results

### The P-body scaffold protein EDC4 is a *bona fide* ATG8 interactor in *Marchantia polymorpha*

To determine whether ribonucleoprotein (RNP) granules are targeted for autophagic degradation in plants, we generated *Marchantia polymorpha* and *Arabidopsis thaliana* lines expressing GFP-tagged Decapping Protein 1 (DCP1), a core component of P-bodies^28^. These lines were established in both wild-type backgrounds (Tak-1 for Marchantia, Col-0 for Arabidopsis) and autophagy-deficient mutants (*atg7-1* and *atg5-1*, respectively). We then performed autophagic flux assays using Torin-1, an mTOR inhibitor that robustly induces autophagy, followed by recovery in the presence or absence of vacuolar V-ATPase or protease inhibitors^29^. In wild-type backgrounds, both MpDCP1 and AtDCP1 underwent vacuolar degradation, whereas this degradation was abolished in *atg7-1* and *atg5-1* mutants (Fig. 1A, Fig. S1A, Fig. S2A–B). We next assessed whether DCP1-positive puncta co-localized with autophagosomes. In Arabidopsis, DCP1 co-localized with ATG8 only upon treatment with the vacuolar V-ATPase inhibitor concanamycin A (Fig. S1B). In contrast, in Marchantia, DCP1 co-localized with ATG8 puncta even under untreated conditions (Fig. 1B). We further found that MpDCP2, another core P-body component, was also degraded via autophagy in Tak-1 but remained stable in the *atg7-1* mutant (Fig. S1C, Fig. S2C–D). Together, these results demonstrate that P-bodies are selectively turned over by autophagy in plants.

**Figure 1.**
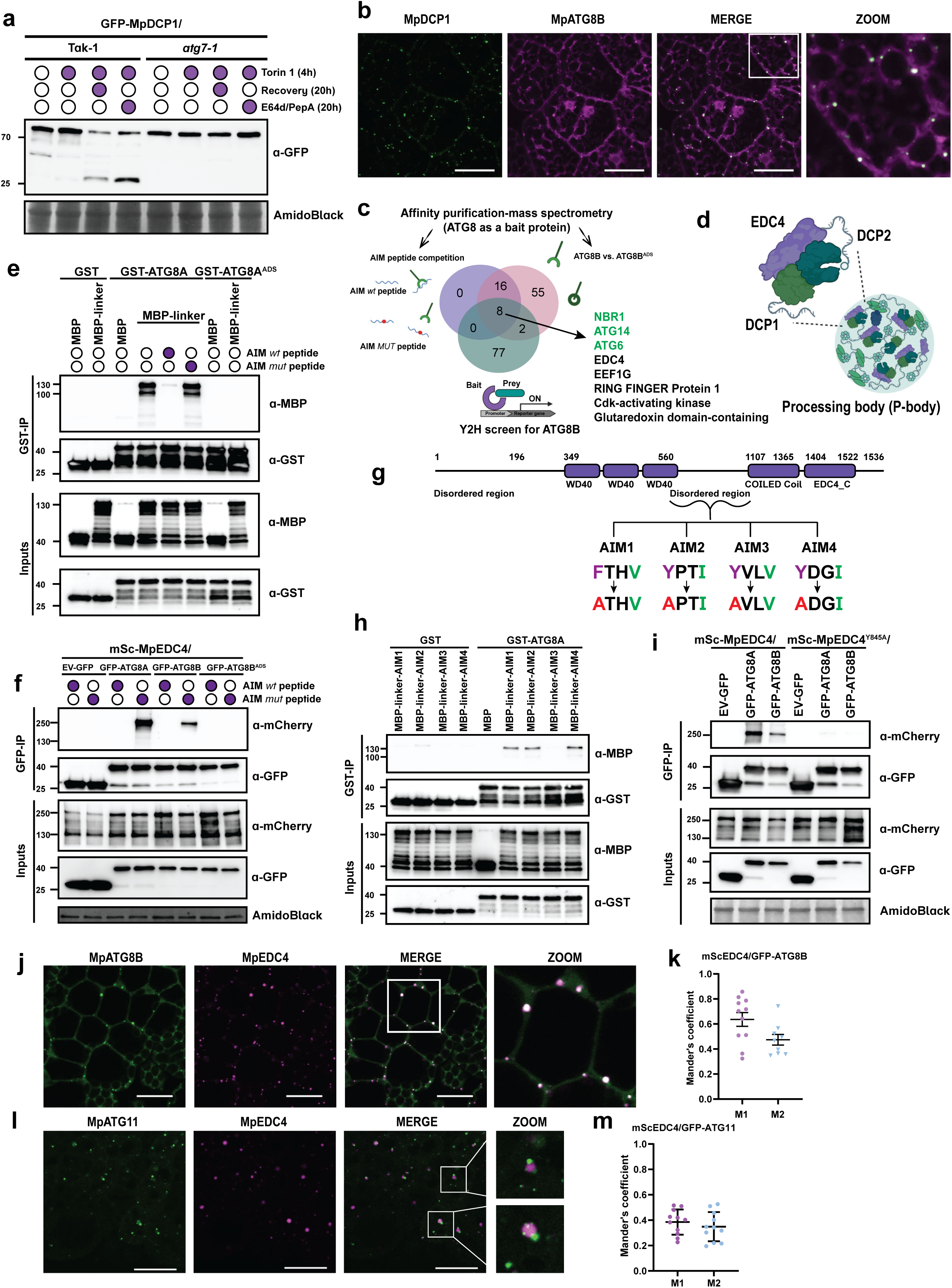
The P-body scaffold protein EDC4 is a *bona fide* ATG8 interactor in *Marchantia polymorpha*. **a, MpDCP1 undergoes vacuolar degradation in an autophagy-dependent manner.** Western blot showing GFP-MpDCP1 flux assay in Tak-1 or *atg7-1* mutant. 6-days old *M. polymorpha* gemmae were incubated in either DMSO or 12 μM Torin-1-containing media (0.5 Gamborg, MES, 1% sucrose) for 4 h, after which plants were either harvested or recovered for additional 20 h with either fresh media (Rec) or media containing 10 μM E64d and 10 μM pepstatin A (E64d/PepA) before harversing. Total protein levels were analyzed by staining with AmidoBlack. A second independent biological replicate is shown in Fig. S2A. **b, MpDCP1 co-localizes with MpATG8-decorated autophagosomes**. Confocal microscopy images of 2-days old *M. polymorpha* thallus cells co-expressing GFP-MpDCP1 with mScarlet-MpATG8B. Thalli were incubated in 0.5 Gamborg, MES, 1% sucrose media. Representative images of minimum 10 biological replicates are shown. Zoom panels show enlarged sections of the white-boxed areas highlighted in the merge panel. Scale bar, 20 μm. **c, EDC4 interacts with ATG8.** Venn diagram comparing the interactomes identified from Yeast-Two-Hybrid and peptide-competition coupled affinity purification mass spectrometry. Data from Yeast Two-Hybrid screen represented in the green circle, data from AP-MS for MpATG8B-associated and MpATG8B^ADS^ sensitive proteins in pink circle, and MpATG8B peptide-outcompeted proteins in violet circle. **d, MpEDC4 is a part of the P-bodies.** EDC4 acts a scaffold protein for decapping proteins 1 and 2, DCP1 and DCP2, respectively. **e, MpEDC4 linker region associates with MpATG8A in AIM-dependent manner *in vitro*.** *In-vitro* pulldown assay coupled to peptide competition showing the association between MpEDC4 linker and MpATG8A. Bacterial lysates containing recombinant protein were mixed and pulled down with GST magnetic agarose beads. Input and bound proteins were immunoblotted with anti-GST and anti-MBP antibodies. AIM *wt* and AIM *mut* peptides were added to the final lysates at a concentration of 200 μM. A second biological replicate is shown in Fig. S2E. ATG8A^ADS^ = ATG8A^(Y52A,L53A)^. **f, MpEDC4 associates with MpATG8A and MpATG8B in an AIM-dependent manner.** *In-planta* GFP-Trap Co-IP coupled to peptide competition showing the association between mScarlet-tagged MpEDC4 and GFP-tagged MpATG8A and MpATG8B isoforms. 9-days old M.polymorpha gemmae were incubated in 0.5 Gamborg, MES, 1% sucrose prior to harvesting. AIM *wt* and AIM *mut* peptides were added to the final lysates at the concentration of 200 μM. Protein extracts were immunoblotted with anti-GFP and anti-mCherry antibodies. A second biological replicate is shown in Fig. S2F. ATG8B^ADS^ = ATG8B^(K51A)^. **g, MpEDC4 harbours 4 putative AIM motifs within its disordered region.** Schematic domain architecture of MpEDC4 with highlighted putative AIMs within the disordered region. AIM residues and respective mutations are shown. AIM1 = EDC4^(F718A)^, AIM2 = EDC4^(Y725A)^, AIM3 = EDC4^(Y845A)^, AIM4 = EDC4^(Y951A)^. **h, AIM3 is the functional AIM of EDC4.** Site-directed mutagenesis of putative AIMs coupled to *in-vitro* pulldown assay reveals AIM3 is the functional AIM. Bacterial lysates containing recombinant protein were mixed and pulled down with GST magnetic agarose beads. Input and bound proteins were immunoblotted with anti-GST and anti-MBP antibodies. A second biological replicate is shown in Fig. S2G. **i, Validation of AIM3 *in vivo*.** *In-planta* GFP-Trap Co-IP showing that mutation of AIM3 (EDC4^(Y845A)^) is sufficient to disrupt association of EDC4 with ATG8. 9-days old *M. polymorpha* gemmae were incubated in 0.5 Gamborg, MES, 1% sucrose prior to harvesting. Protein extracts were immunoblotted with anti-GFP and anti-mCherry antibodies. aimMUT = EDC4^(Y845A)^. A second biological replicate is shown in Fig. S2H. **j-k, MpEDC4 co-localizes with MpATG8-decorated autophagosomes.** Confocal microscopy images of 2-days old *M. polymorpha* thallus cells co-expressing GFP-MpATG8B with mScarlet-MpEDC4. Thalli were incubated in 0.5 Gamborg, MES, 1% sucrose media. Representative images of minimum 10 biological replicates are shown. Zoom panels show enlarged sections of the white-boxed areas highlighted in the merge panel. Scale bar, 20 μm. Quantification of confocal micrographs in Fig. 1J as assessed by the Mander’s colocalization coefficients M1 and M2. Bars indicate the mean ± SD of minimum 10 biological replicates. **l-m, MpATG11 forms initiation hubs on the surface of EDC4-positive puncta.** Confocal microscopy images of 2-days old *M. polymorpha* thallus cells co-expressing GFP-MpATG11 with mScarlet-MpEDC4. Thalli were incubated in 0.5 Gamborg, MES, 1% sucrose media. Representative images of minimum 10 biological replicates are shown. Zoom panels show enlarged sections of the white-boxed areas highlighted in the merge panel. Scale bar, 20 μm. Quantification of confocal micrographs in Fig. 1J as assessed by the Mander’s colocalization coefficients M1 and M2. Bars indicate the mean ± SD of minimum 10 biological replicates.

To identify selective autophagy receptors (SARs) that mediate P-body turnover, we performed ATG8 affinity purification coupled to mass spectrometry (AP-MS) in *Marchantia polymorpha*. To enrich for AIM-dependent ATG8 interactors, the AP-MS was conducted in the presence of an AIM peptide, which selectively competes for ATG8 binding and facilitates the identification of direct AIM-dependent interactors^30,31^. In parallel, we performed AP-MS using an ATG8^ADS^ mutant that is defective in AIM binding^32^. This combined approach yielded 24 candidate proteins whose interaction with ATG8 was disrupted both by AIM peptide competition and lost in the ATG8^ADS^ background. To further refine our candidate list, we carried out a genome-wide yeast two-hybrid (Y2H) screen using MpATG8A and MpATG8B as baits. Eight proteins overlapped between the AP-MS and Y2H datasets (Fig. 1C, Supplementary Data 1), including three known ATG8 interactors—NBR1, ATG14, and ATG6—thereby validating our approach^33,34^. Notably, among the remaining five candidates was Enhancer of mRNA Decapping 4 (EDC4), also known as Varicose (VCS) or Ge-1, a highly conserved P-body scaffold protein known to interact with DCP1 and DCP2^35^ (Fig. 1D).

We next investigated the interaction between EDC4 and ATG8 using *in vitro* pulldown assays. Although full-length MpEDC4 could not be expressed in *E. coli*, we focused on the region between the terminal WD40 domain and the predicted coiled-coil domain, which had been identified as a potential ATG8B-interacting region in the Y2H screen (Fig. 1G). This linker region of MpEDC4 bound MpATG8A in an AIM-dependent manner, as the interaction was abolished by addition of the AIM wild-type peptide and was lost when using the ATG8^ADS^ mutant (Fig. 1E, Fig. S2E). To assess whether EDC4 also interacts with ATG8 *in vivo*, we generated *Marchantia polymorpha* lines expressing mScarlet-tagged EDC4 in the GFP- ATG8A and GFP-ATG8B backgrounds. GFP alone and GFP-MpATG8B^ADS^ were used as negative controls. Co-immunoprecipitation (Co-IP) experiments revealed that MpEDC4 associates with both ATG8 isoforms in a manner sensitive to AIM peptide competition, and this interaction was not observed with GFP alone or GFP-MpATG8B^ADS^ (Fig. 1F, Fig. S2F).

To pinpoint the functional ATG8-interacting motif (AIM) within EDC4, we analyzed its linker region using the iLIR tool (https://ilir.warwick.ac.uk), which predicted four candidate AIMs (Fig. 1G). We individually mutated the conserved aromatic residue in each AIM to alanine and assessed ATG8 binding via *in vitro* pulldown assays. Substitution of tyrosine 845 with alanine in the third predicted motif (EDC4^Y845A^) was sufficient to abolish the interaction between the EDC4 linker region and MpATG8A (Fig. 1H, Fig. S2G). To validate this interaction *in vivo*, we generated *Marchantia polymorpha* lines expressing either mScarlet tagged wild-type EDC4 or Y845A mutant in GFP-ATG8A and GFP-ATG8B backgrounds. Co-immunoprecipitation experiments revealed that this single amino acid substitution in AIM3 nearly abolished the interaction between EDC4 and ATG8 (Fig. 1I, Fig. S2H). These findings demonstrate that MpEDC4 engages ATG8 via a functional AIM centered on tyrosine 845.

We next investigated whether EDC4 co-localizes with key autophagy components ATG8 and ATG11^36,37^. EDC4 and ATG8 displayed nearly complete spatial overlap, indicating that EDC4-positive puncta are decorated by ATG8 (Fig. 1J–K). To exclude the possibility that this co-localization results from EDC4 overexpression under the constitutive *EF1α* promoter, we generated transgenic lines expressing GFP-tagged MpEDC4 under its native promoter (*pEDC4::GFP-MpEDC4*) in an *mScarlet-ATG8B* background. Similar co-localization patterns were observed, confirming that the interaction is not an artefact of overexpression (Fig. S1D). In contrast, ATG11 localized to the periphery of EDC4-positive puncta, forming ring-like structures around the granules (Fig. 1L–M). This peripheral localization of ATG11 is consistent with its proposed role in establishing initiation hubs on selective cargo to trigger autophagosome formation5. Together, these findings show that EDC4 co-localizes with both ATG8 and ATG11, further supporting its function as an autophagy substrate and a potential cargo receptor for selective autophagy.

### MpEDC4 is a conserved P-body component with a lineage-specific AIM

We next asked whether *Marchantia polymorpha* EDC4 functions as a bona fide P-body component. Phylogenetic analysis revealed that MpEDC4 is related to ancestral EDC4 orthologs, including the well-characterized human and Arabidopsis homologs, suggesting strong evolutionary conservation (Fig. S3A–B). Interestingly, sequence alignment showed that while the overall domain architecture of EDC4 is conserved across eukaryotes, the functional AIM motif is only conserved within the Marchantiaceae family (Fig. 2A, Fig. S3C). This suggests that MpEDC4 acquired an ATG8-interacting motif, linking it to the autophagy pathway. Consistently, the Arabidopsis EDC4 homolog VCS bound neither ATG8A nor ATG8E in *in vitro* pulldown assays (Fig. S3D).

**Figure 2.**
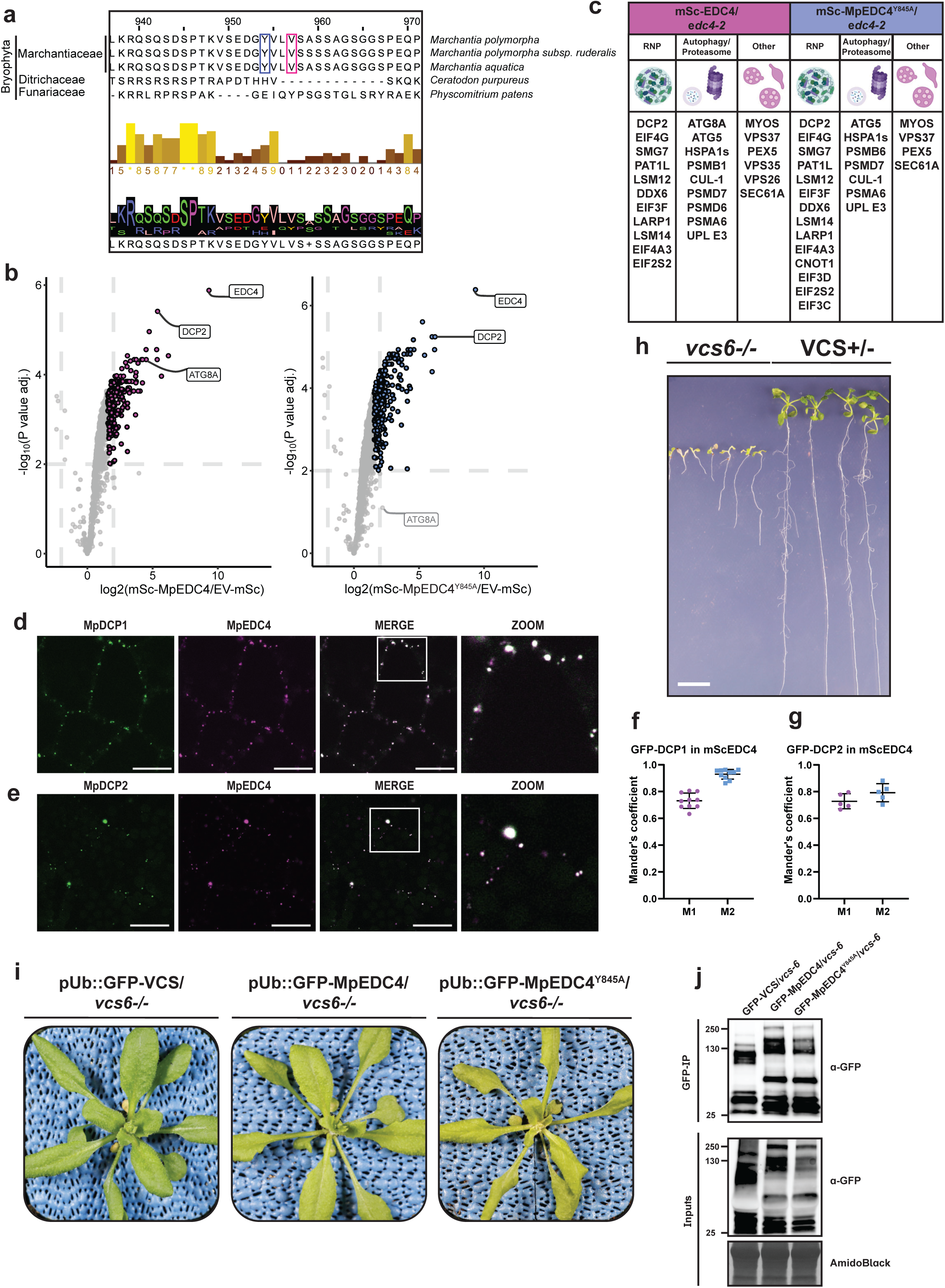
MpEDC4 is a conserved P-body component with a lineage-specific AIM. **a, Multiple sequence alignment of EDC4 in Bryophyta revealed that only Marchantiaceae EDC4s has a canonical AIM motif within the disordered region. b, Volcano plots representing *in-planta* RFP-trap Co-IP coupled to AP-MS showing enrichment of proteins co-purified with mScarlet-tagged MpEDC4 or MpEDC4^Y845A^ compared to mScarlet alone control.** 9-days old *M.polymorpha* gemmae were incubated in 0.5 Gamborg, MES, 1% sucrose prior to harvesting. The horizontal dashed line indicates the threshold above which proteins are significantly enriched (p value adjusted < 0.05) and the vertical dashed line indicated the threshold for which proteins log_2_ fold change is above 2. **c, MpEDC4 associates with markers of P-bodies.** Top ranked candidates from AP-MS were distributed into 3 groups: canonical P-bodies markers, proteins linked to autophagy and proteasomal degradation and others. **d-g, MpEDC4 co-localizes with MpDCP1 and MpDCP2.** Confocal microscopy images of 2-days old *M. polymorpha* thallus cells co-expressing mScarlet-MpEDC4 with either GFP-MpDCP1 or GFP-MpDCP2. Thalli were incubated in 0.5 Gamborg, MES, 1% sucrose media. Representative images of minimum 10 biological replicates are shown. Zoom panels show enlarged sections of the white-boxed areas highlighted in the merge panel. Scale bar, 20 μm. Quantification of confocal micrographs in Fig. 2C and Fig. 2E as assessed by the Mander’s colocalization coefficients M1 and M2. Bars indicate the mean ± SD of minimum 10 biological replicates. **h, AtVCS mutant *vcs-/-* (*vcs-6),* has severe developmental defects, leading to seedling lethality**. 10-days old Arabidopsis seedlings of segregating *VCS+/-* heterozygotes were grown on 1% agar plates supplemented with ½ MS, MES and 1% Sucrose. After 10 days, seedlings were separated, *vcs-6* are shown on the left part of the plate, VCS *wt* and *VCS+/-* heterozygotes are shown on the right. Scale bar, 1 cm. **i, MpEDC4 rescues Arabidopsis *vcs-6* mutant phenotype.** Complementation of *VCS+/-* heterozygotes with GFP-tagged AtVCS, MpEDC4 or MpEDC4^Y845A^ rescues *vcs-6* plants in the next generation. Approximately 3-weeks old Arabidopsis plants were imaged growing on soil. **j, *In planta* GFP-Trap Co-IP showing the expression of GFP-tagged AtVCS in *vcs-6*, as well as GFP-tagged MpEDC4 *wt* or MpEDC4^AIM-^ in *vcs-6*.** 10-days old *A. thaliana* seedlings were incubated in ½ MS, MES, 1% sucrose prior to harvesting. Protein extracts were immunoblotted with anti-GFP antibodies.

To further investigate MpEDC4 function, we performed affinity purification followed by mass spectrometry (AP-MS) using mScarlet-tagged wild-type EDC4 and the AIM-disrupted EDC4^Y845A^ variant; mScarlet alone served as a negative control. As expected, ATG8 was among the top interactors in wild-type EDC4 pulldowns. Surprisingly, ATG8 was also recovered, albeit at reduced levels, in the EDC4^Y845A^ interactome, suggesting the presence of additional adaptor proteins that may mediate P-body autophagy. ATG5 was also enriched, consistent with EDC4 being targeted via canonical autophagy pathways (Fig. 2B).

Both wild-type and AIM-mutant EDC4 co-purified with established P-body components including DCP2, Pat1, DDX6, translation initiation factors, and members of the CCR4-NOT complex, reinforcing its conserved role as a P-body scaffold^38^. In addition to these canonical factors, we identified proteasome subunits and E3 ligases, pointing to a potential role for the ubiquitin–proteasome system in P-body turnover, as recently suggested for other cytoplasmic granules such as stress granules in human and Arabidopsis studies^9,39^. Notably, we also detected retromer complex components (VPS26 and VPS35), which are involved in endosome-to-Golgi and plasma membrane traffickings (Fig. 2B–C). Apart from retromer complex components, we also detected VPS37, which is canonically associated with the ESCRT-I complex^40^, hinting at possible coordination between degradative and recycling pathways

To confirm EDC4 subcellular localization, we generated stable Marchantia lines co-expressing mScarlet-tagged EDC4 and GFP-tagged DCP1 or DCP2. Confocal imaging revealed strong co-localization between EDC4 and both decapping factors, consistent with P-body localization (Fig. 2D–G).

Finally, to assess functional conservation in mRNA decapping, we performed cross-species complementation in Arabidopsis. The *vcs-/- (vcs-6)* mutant displays severe developmental defects (Fig. 2H), including stunted growth and failure to form true leaves^41^. Ectopic expression of either Arabidopsis EDC4, Marchantia EDC4, or the AIM-deficient EDC4^Y845A^ fully rescued the mutant phenotype, indicating that MpEDC4 retains its ancestral function as a P-body scaffold (Fig. 2I-J).

Taken together, these findings support a model in which a conserved P-body protein in Marchantiaceae has acquired an AIM motif, enabling its integration into the autophagy pathway while preserving its ancestral role in RNA metabolism.

### Autophagic flux analysis suggest multiple pathways mediate P-body recycling

To investigate the degradation dynamics of MpEDC4, we first performed triple co-localization analysis using fluorescently labeled MpEDC4, MpDCP1, and MpATG8A. The strong spatial overlap among all three proteins suggests that MpEDC4 is recruited to autophagosomes as part of P-body complexes (Fig. 3A).

**Figure 3.**
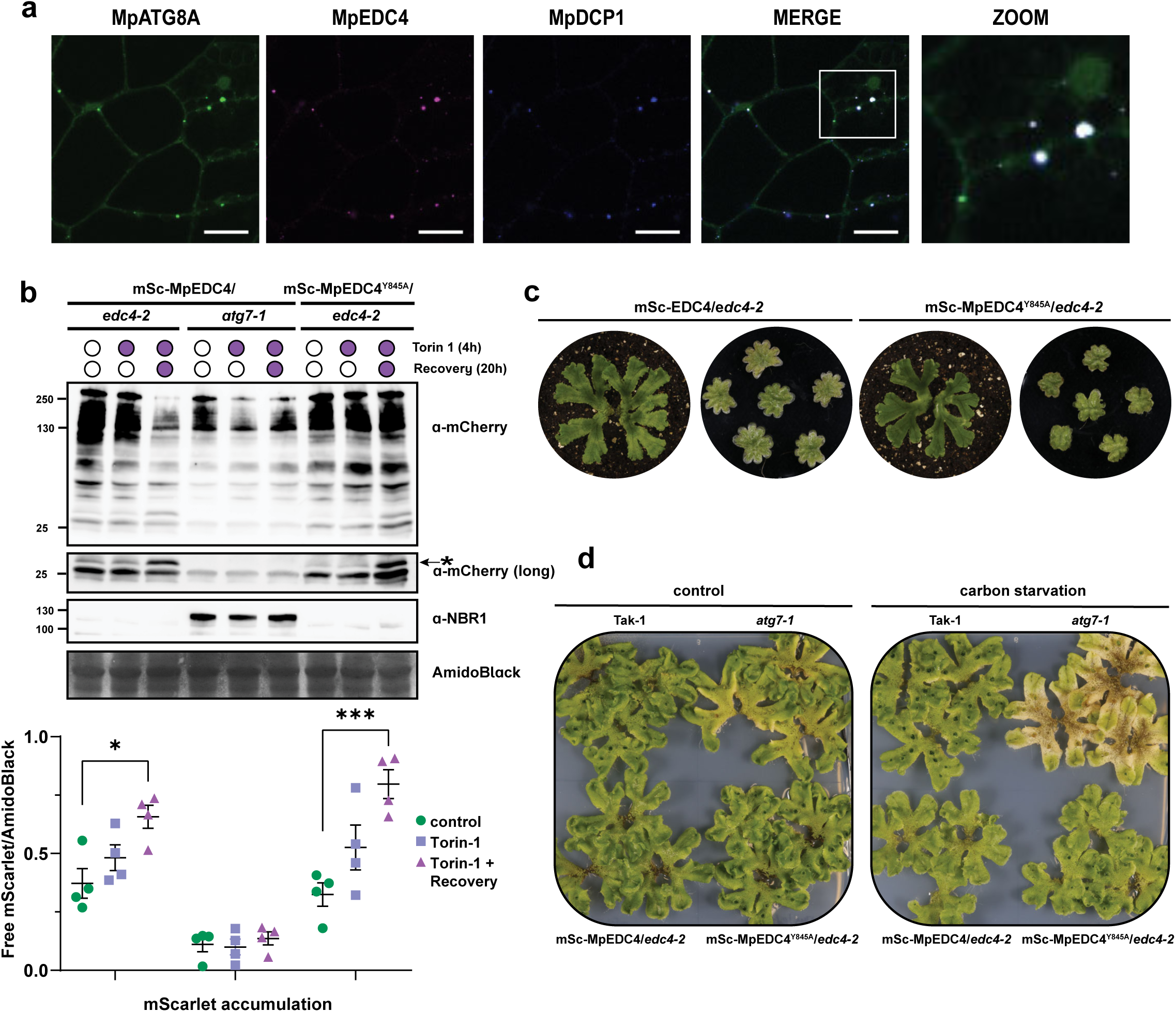
Autophagic flux analysis reveals multiple pathways mediate EDC4 turnover. **a**, **Core P-body proteins, MpEDC4 and MpDCP1, co-localize with MpATG8-decorated autophagosomes.** Confocal microscopy images of 2-days old *M. polymorpha* thallus cells co-expressing GFP-MpATG8B with mSc-MpEDC4 and MpDCP1-mKalama-3xflag. Thalli were incubated in 0.5 Gamborg, MES, 1% sucrose media. Representative images of minimum 10 biological replicates are shown. Zoom panels show enlarged sections of the white-boxed areas highlighted in the merge panel. Scale bar, 20 μm. **b, MpEDC4^Y845A^ undergoes vacuolar degradation in *edc4-/-* plants (*edc4-2*).** Western blot showing mSc-MpEDC4 *wt* and mSc- MpEDC4^Y845A^ flux assay and endogenous NBR1 level in *edc4-2* mutant background. Plants expressing mScarlet-tagged MpEDC4 *wt* in *atg7-1* mutant were used as negative controls. 6- days old *M. polymorpha* gemmae were incubated in either DMSO or 12 μM Torin-1-containing media (0.5 Gamborg, MES, 1% sucrose) for 4 h, after which plants were either harvested or recovered for additional 20 h with fresh media (Rec) before harvesting. Total protein level was analyzed by staining with AmidoBlack. Three independent biological replicates are shown in Fig. S4B-D. *: free mScarlet band. **c, Marchantia expressing mScarlet-tagged MpEDC4^Y845A^ in *edc4-2* background show no phenotype compared to plants expressing *wt* protein.** Approximately, 3 weeks *M. polymorpha* gemmae expressing either mScarlet- MpEDC4 *wt* or MpEDC4**^Y845A^** in *edc4-2* mutant background were grown under far-red-enriched irradiation (left side) followed by another 3 weeks until antheridiophores were mature. Mature antheridiophores were cut and imaged (right side). **d, Plants expressing MpEDC4^Y845A^ is insensitive to carbon starvation.** 12-days old *M. polymorpha* gemmae were transferred to the 1% agar plates supplemented with 0.5 Gamborg and MES, without sucrose, followed by transferring to either normal light irradiation or in the darkness for 2 days. *atg7-1* mutant plants were used as control. A second independent biological replicate is shown in Fig. S5D.

We next hypothesized that MpEDC4^Y845A^, which lacks a functional AIM, would not undergo autophagic degradation. To test this, we generated *edc4* CRISPR-Cas9 knockout lines expressing either mScarlet-MpEDC4^WT^ or mScarlet-MpEDC4^Y845A^ (Fig. S4A). As a positive control for autophagy inhibition, we used *atg7-1* mutants expressing mScarlet-MpEDC4. As expected, no free mScarlet fragment—an indicator of autophagic flux—was detected in the *atg7-1* background. However, both mSc-MpEDC4^WT^/*edc4-2* and mSc-MpEDC4^Y845A^/*edc4-2* lines displayed detectable levels of free mScarlet, indicating that MpEDC4 can still undergo autophagic degradation independently of ATG8 binding (Fig. 3B, Fig. S4B–D).

To rule out the possibility that this degradation was due to overexpression under the constitutive EF1α promoter, we used a transgenic line expressing GFP-tagged MpEDC4 under its native minimal promoter (pEDC4::GFP-MpEDC4). Treatment with the vacuolar protease inhibitors E64d and pepstatin A led to the accumulation of full-length GFP-MpEDC4, confirming that MpEDC4 is degraded via the vacuole and that this process is not an artifact of overexpression (Fig. S4E).

To further validate these findings, we generated independent *edc4* knockout lines using alternative guide RNAs (mSc-MpEDC4^WT^/*edc4-1* and mSc-MpEDC4^Y845A^/*edc4-1*) and induced autophagy using salt stress. Similar to Torin-1 treatment, we observed free mScarlet fragments in both backgrounds, supporting the involvement of an alternative degradation pathway (Fig. S5A–C).

We next assessed whether expression of MpEDC4^Y845A^ causes any physiological defects. Plants expressing either wild-type or AIM-mutant MpEDC4 were morphologically indistinguishable (Fig. 3C). Previous work showed that autophagy is critical for spermatozoid maturation in Marchantia^42,43^. Both lines produced morphologically normal antheridiophores and functional spermatozoids, indicating that EDC4 AIM function is dispensable for this developmental process (Fig. 3C, Fig. S4F). Similarly, in carbon starvation assays, only *atg7- 1* mutants exhibited early senescence, suggesting that EDC4 is not a general regulator of autophagy (Fig. 3D, Fig. S5D).

Given previous evidence implicating the role of ubiquitination in the disassembly of membraneless RNP granules^9,39^, we also examined whether the ubiquitin-binding aggrephagy receptor NBR1^21^ could participate in EDC4 turnover. We only observed a partial co-localization between NBR1 and EDC4/DCP1-positive puncta (Fig. S4G), suggesting the presence of additional, possibly redundant, selective autophagy mechanisms in P-body recycling.

### MpDCP1 interacts with ATG8 via a non-canonical reverse AIM motif

To search for the redundant selective autophagy receptors that could mediate P-body recycling, we revisited our ATG8 interactome datasets^31^ and discovered MpDCP1, another well-known P-body protein^28^ (Fig. 4A), prompting us to test whether MpDCP1 physically interacts with ATG8. *In vitro* pulldown assays confirmed that MpDCP1, like MpEDC4, binds ATG8 in an AIM-dependent manner (Fig. 4B, Fig. S6A).

**Figure 4.**
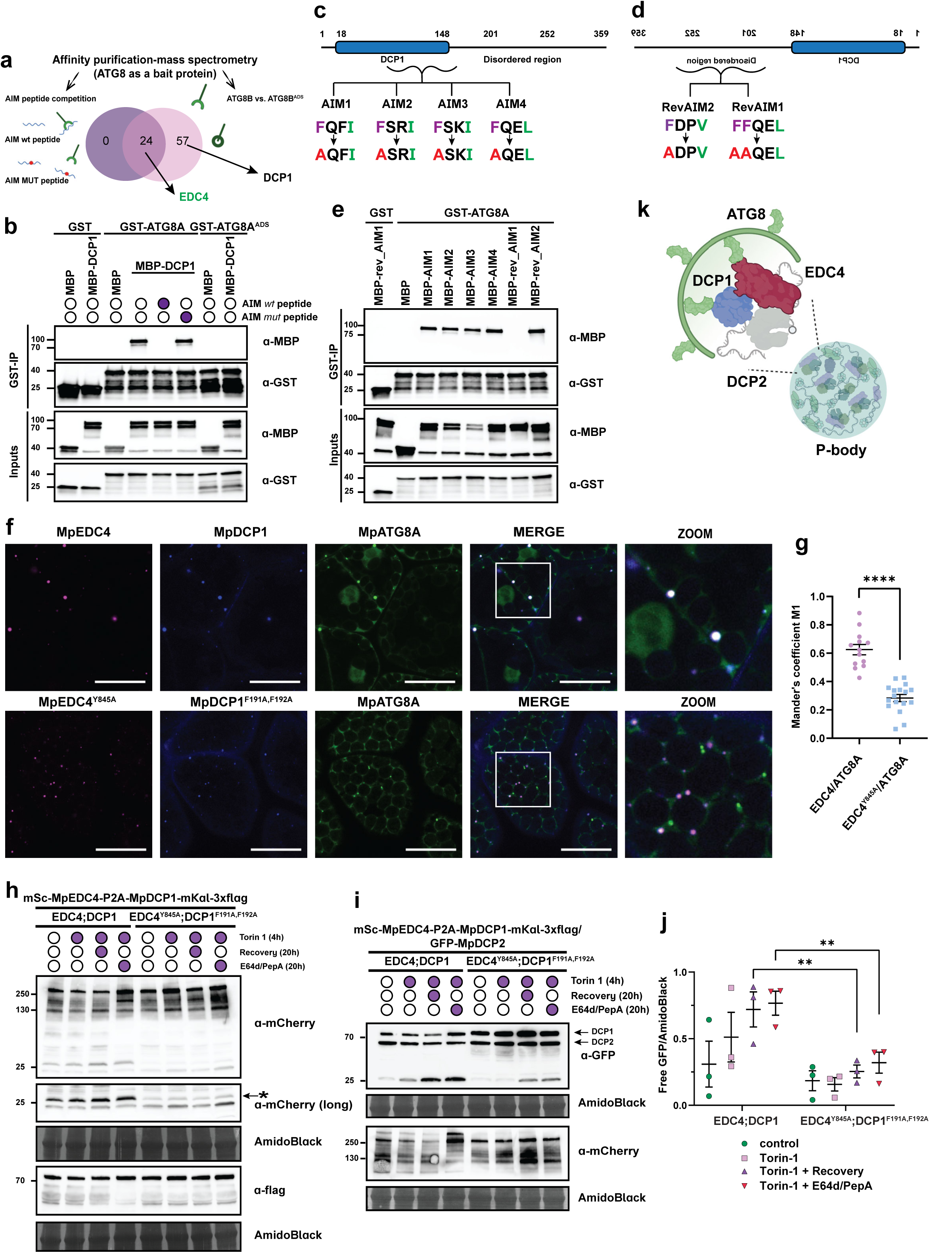
EDC4-DCP1 receptor pair mediate P-body turnover in *Marchantia polymorpha*. **a, DCP1 associates with ATG8.** Venn diagram comparing MpATG8B interactors. MpATG8B^ADS^ sensitive proteins represented in pink circle, peptide-outcompeted proteins in violet circle. **b, MpDCP1 interacts with MpATG8A in AIM-dependent manner.** *In vitro* pulldown assay coupled to peptide competition showing the interaction between MpDCP1 and MpATG8A. Bacterial lysates containing recombinant protein were mixed and pulled down with GST magnetic agarose beads. Input and bound proteins were immunoblotted with anti-GST and anti-MBP antibodies. AIM wt and AIM mut peptides were added to the final lysates at the concentration of 200 μM. A second biological replicate is shown in Fig. S6A. ATG8A^ADS^ = ATG8A^(Y52A,L53A)^. **c, MpDCP1 has 4 putative AIM motifs within its forward sequence.** Schematic domain architecture of MpDCP1 with highlighted putative AIMs. AIM residues and respective mutations are shown. AIM1 = DCP1^(F73A)^, AIM2 = DCP1 ^(F130A)^, AIM3 = DCP1 ^(F137A)^, AIM4 = DCP1 ^(F154A)^. **d, MpDCP1 has 2 putative AIM motifs within its reverse sequence in disordered region.** Schematic domain architecture of MpDCP1 with highlighted putative AIMs in reverse sequence. AIM residues and respective mutations are shown. Rev AIM1 = DCP1^(F191A,F192A)^, RevAIM2 = DCP1 ^(F263A)^. **e, RevAIM1 is a functional AIM.** Site-directed mutagenesis of putative AIMs and RevAIMs coupled to *in-vitro* pulldown assay reveals RevAIM1 is the functional AIM within DCP1 disordered region. Bacterial lysates containing recombinant protein were mixed and pulled down with GST magnetic agarose beads. Input and bound proteins were immunoblotted with anti-GST and anti-MBP antibodies. A second biological replicate is shown in Fig. S6B. **f, MpATG8-decorated autophagosomes co- localize with *wt* autophagy receptors MpEDC4 and MpDCP1 but not with AIM-mutant versions of EDC4 and DCP1.** Confocal microscopy images of 2-days old *M. polymorpha* thallus cells co-expressing GFP-MpATG8B with either *wt* (mSc-EDC4-P2A-DCP1-Kal-3xflag) or *AIM mutant* (mSc-EDC4^Y845A^-P2A-DCP1^F191A,F192A^-Kal1-3xflag). Thalli were incubated in 0.5 Gamborg, MES, 1% sucrose media. Representative images of minimum 10 biological replicates are shown. Zoom panels show enlarged sections of the white-boxed areas highlighted in the merge panel. Scale bar, 20 μm. **g, Quantification of confocal micrographs in** Fig. 4D **as assessed by the Mander’s colocalization coefficients M1 between mScarlet-EDC4 and GFP-ATG8A.** Bars indicate the mean ± SD of minimum 10 biological replicates. Two-tailed Mann-Whitney test was performed to analyze the significance of the differences of M1 values between mScarlet and GFP for double *wt* and double *AIM*. ****, P value < 0.0001. **h, MpEDC4 undergoes vacuolar degradation in an EDC4-DCP1- dependent manner.** Western blot showing mScarlet-MpEDC4 flux assay in either double *wt* (mSc-EDC4-P2A-DCP1-Kal-3xflag) or *aimMUT* (mSc-EDC4^Y845A^-P2A-DCP1^F191A,F192A^-Kal1-3xflag) plants. 6-days old *M.polymorpha* gemmae were incubated in either DMSO or 12 μM Torin-1-containing media (0.5 Gamborg, MES, 1% sucrose) for 4 h, after which plants were either harvested or recovered for additional 20 h with either fresh medium (Rec) before harvesting. Total protein level was analyzed by staining with AmidoBlack. Second independent biological replicate is shown in Fig. S6C. *:free mScarlet band. **I-j, MpDCP2 undergoes vacuolar degradation in an EDC4-DCP1-dependent manner.** Western blot showing GFP- MpDCP2 flux assay in either double *wt* (mSc-EDC4-P2A-DCP1-Kal-3xflag) or *aimMUT* (mSc- EDC4^Y845A^-P2A-DCP1^F191A,F192A^-Kal1-3xflag) plants. 6-days old *M. polymorpha* gemmae were incubated in either DMSO or 12 μM Torin-1-containing media (0.5 Gamborg, MES, 1% sucrose) for 4 h, after which plants were either harvested or recovered for additional 20 h with either fresh media (Rec) or media containing 10 μM E64d and 10 μM pepstatin A (E64d/PepA) before harvesting. Total protein levels were analyzed by staining with AmidoBlack. Two independent biological replicates are shown in Fig. S6D-E. **k, Proposed model for selective autophagic degradation of P-bodies via the EDC4-DCP1 receptor pair.**

To identify the underlying motif mediating this interaction, we analyzed the MpDCP1 sequence using the iLIR prediction tool (https://ilir.warwick.ac.uk), which identified four candidate AIMs (Fig. 4C). However, alanine substitutions within these motifs failed to disrupt binding, suggesting that the canonical predicted AIMs were non-functional (Fig. 4E). Given these results, we hypothesized that MpDCP1 might contain a non-canonical AIM. Re-analysis of the sequence in reverse orientation revealed several putative reverse AIMs, including two motifs located within the protein’s intrinsically disordered region (IDR) (Fig. 4D). Mutation of two conserved phenylalanines (F191 and F192) within reverse AIM1 to alanines (MpDCP1^F191A,F192A^) abolished ATG8A binding *in vitro*, confirming the functional importance of this non-canonical motif (Fig. 4E, Fig. S6B). These results establish that MpDCP1 interacts with ATG8 via a previously uncharacterized reverse AIM motif.

We next examined whether MpEDC4 and MpDCP1 function together to mediate P-body recycling. To test this, we co-expressed wild-type or AIM-disrupted versions of both proteins in the GFP-MpATG8A background using self-cleaving P2A constructs encoding either wild-type (mSc-EDC4–P2A–DCP1–mKal–3×FLAG) or mutant (mSc-EDC4^Y845A^–P2A– DCP1^F191A,F192A^–mKal–3×FLAG) variants. In wild-type lines, we observed strong co- localization between MpEDC4, MpDCP1, and ATG8A (Fig. 4F). In contrast, in the double AIM mutant line, EDC4–DCP1 puncta were spatially separated from ATG8A, indicating a loss of autophagic targeting (Fig. 4G).

To determine whether this impaired localization translates into reduced degradation, we performed autophagic flux assays following Torin-1 treatment. Double AIM mutant lines exhibited markedly reduced cleavage of mSc-MpEDC4 compared to wild-type, indicating that both DCP1 and EDC4 contribute to autophagy-mediated turnover of EDC4 (Fig. 4H, Fig. S6C). We extended this approach to lines expressing the same constructs in a GFP-MpDCP2 background. Again, we observed reduced GFP cleavage in the AIM mutant compared to wild-type, suggesting that both MpEDC4 and MpDCP1 are required for the efficient autophagic degradation of DCP2 (Fig. 4I–J, Fig. S6D–E). These findings support a model in which EDC4 and DCP1 act cooperatively to mediate the selective autophagic recycling of P-bodies in *Marchantia polymorpha* (Fig. 4K).

Finally, we asked whether the reverse AIM motif identified in MpDCP1 is conserved across species. Phylogenetic analysis revealed that DCP1 orthologs are present across all major eukaryotic clades (Fig. S7A–B). However, sequence alignment showed that the reverse AIM motif is not conserved beyond Marchantiaceae. *In vitro* pulldown assays confirmed that the Arabidopsis DCP1 ortholog (AtDCP1) does not bind ATG8A, in contrast to the positive control ER-phagy receptor C53 (Fig. S7D, Fig. S6F). These data suggest that the DCP1–ATG8 interaction is specific to Marchantia, supporting the notion that this liverwort evolved a lineage-specific autophagy receptor pair to mediate P-body recycling.

### MpEDC4 expression in human cells promotes α-Synuclein clearance

To explore the potential broader relevance of a lineage-specific autophagy receptor, we asked whether MpEDC4 could function in a heterologous system. A recent study reported that human EDC4 (HsEDC4) binds α-Synuclein, a protein genetically and pathologically linked to Parkinson’s disease (PD)^44^. Given the evolutionary conservation of the autophagy machinery, we hypothesized that MpEDC4 could facilitate α-Synuclein degradation when ectopically expressed in human cells (Fig. 5A).

**Figure 5.**
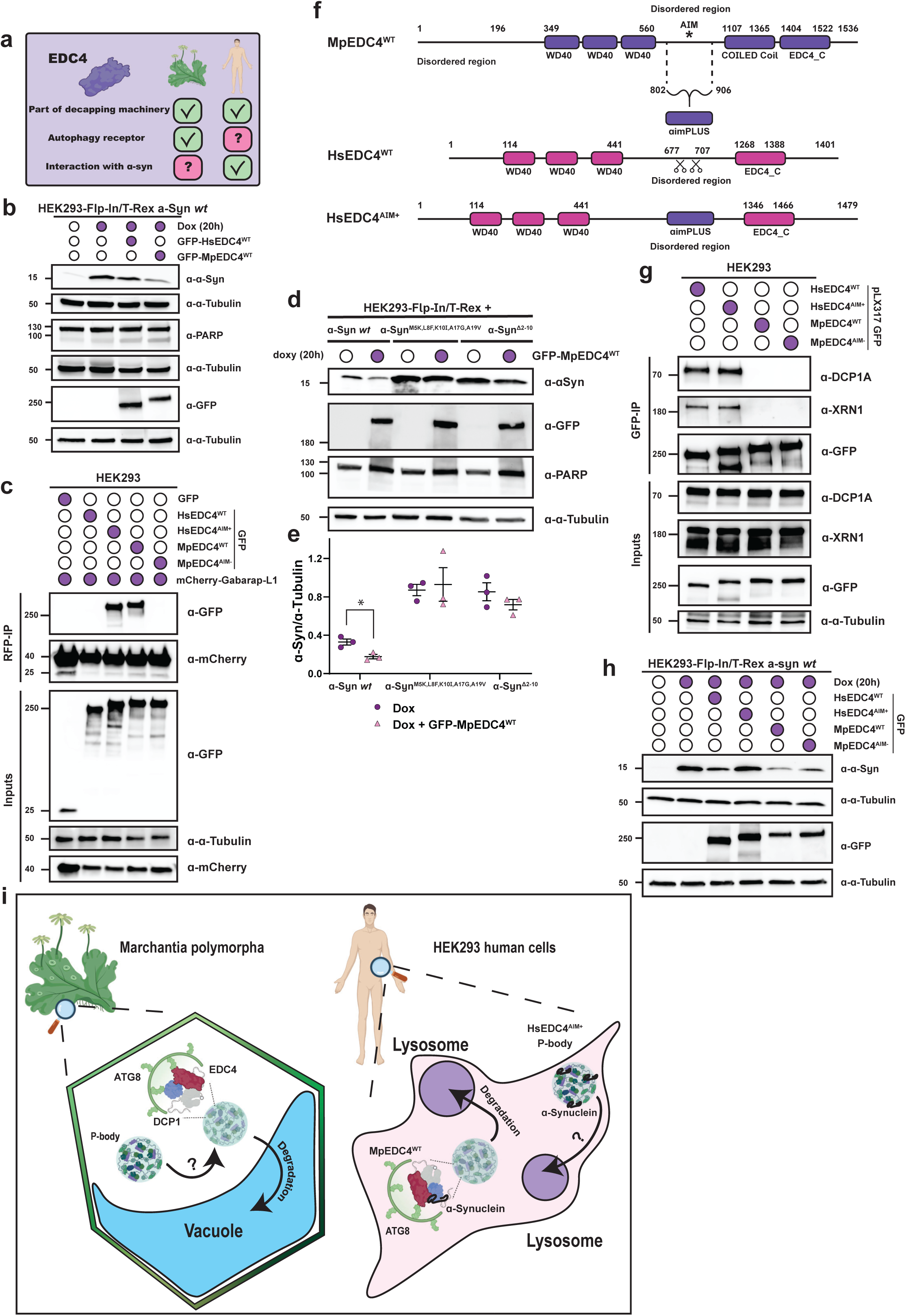
MpEDC4 expression in human cells promotes α-Synuclein clearance. **a, Schematic summary of EDC4 functions in Marchantia and human. b,** MpEDC4 facilitates α-Synuclein clearance in HEK293-Flp-In/T-Rex cells expressing α-Synuclein upon doxycycline treatment. Cells were transfected for 24 hours with GFP-tagged EDC4 (either Hs or Mp), followed by overnight treatment with 250 μg/μl doxycycline. Total protein levels were analyzed by staining with α-tubulin antibodies. A second independent biological replicate is shown in Fig. S8A. **c, Chimeric HsEDC4^AIM+^ associates with HsATG8 ortholog, Gabarap-L1.** *In vivo* RFP-Trap Co-IP in HEK293T cells showing that both GFP-tagged HsEDC4^AIM+^ chimeric protein and MpEDC4^WT^ are associated with mCherry-tagged Gabarap-L1, while HsEDC4^WT^ and MpEDC4^AIM-^ are not. Cells expressing GFP alone were used as negative control. Protein extracts were immunoblotted with anti-GFP and anti-mCherry antibodies. Anti- α-tubulin antibodies were used as loading control for inputs. A second biological replicate is shown in Fig. S8B. **d, α-Synuclein degradation is EDC4-dependent.** WB showing stabilization of α-Synuclein mutants that either cannot bind EDC4 (α- Synuclein^M5K,L8F,K10I,A17G,A19V^) or fails to bind both EDC4 and membranes (α-Synuclein^Δ2-10^). Cells were transfected for 24 hours with GFP-tagged EDC4 followed by overnight treatment with 250 μg/μl doxycycline. Total protein levels were analyzed by staining with α-tubulin antibodies. A second and third independent biological replicates are shown in Fig. S8B-C. **e,** Densitometry for Fig. 5C, Fig. S8C-D. **f, Engineering of a chimeric HsEDC4^AIM+^ that harbors MpEDC4 AIM region.** Schematic illustrating the cloning strategy for generation of chimeric HsEDC4 bearing AIM form MpEDC4. The part of MpEDC4 disordered region between D802 an G906 was cloned into the disordered region of HsEDC4 between P677 and S707. **g, HsEDC4^AIM+^ associates with canonical P-bodies markers, DCP1A and XRN1.** *In vivo* GFP- Trap Co-IP in HEK293T cells showing that both GFP-tagged HsEDC4^AIM+^ and HsEDC4^WT^ associates with canonical P-bodies markers, DCP1A and XRN1, unlike MpEDC4^WT^ and MpEDC4^AIM-^. Protein extracts were immunoblotted with anti-GFP, anti-XNR1, anti-DCP1A antibodies. Anti-α-tubulin antibodies were used as loading control for inputs. A second biological replicate is shown in Fig. S8G. **h, HsEDC4^AIM+^ failed to promote α-Synuclein degradation.** Cells were transfected for 24 hours with GFP-tagged EDC4 (either Hs or Mp), followed by overnight treatment with 250 μg/μl doxycycline. Total protein levels were analyzed by staining with α-tubulin antibodies. A second independent biological replicate is shown in Fig. S8H. **i, Working model for EDC4-DCP1-dependent autophagy of P-bodies.** MpEDC4 and MpDCP1 act as a functional autophagy receptor pair that mediates selective degradation of P-bodies in the early land plant *Marchantia polymorpha.* Expression of MpEDC4 in human cells promoted clearance of α-Synuclein, a protein implicated in Parkinson’s disease.

To test this, we expressed GFP-tagged MpEDC4 or HsEDC4 in HEK293-Flp-In/T-REx cells engineered to express α-Synuclein upon doxycycline induction. Strikingly, expression of MpEDC4 resulted in robust α-Synuclein clearance. In contrast, HsEDC4 had no significant effect (Fig. 5B, Fig. S8A). No PARP cleavage was observed upon MpEDC4 expression, ruling out cytotoxicity. Co-immunoprecipitation (co-IP) experiments further revealed that MpEDC4— but not HsEDC4—interacts with the human ATG8 ortholog GABARAP, suggesting that MpEDC4 engages the autophagy machinery in human cells (Fig. 5C, Fig. S8B).

To determine whether EDC4-dependent degradation of α-Synuclein requires physical interaction, we used α-Synuclein variants with mutations that were reported to disrupt its binding to HsEDC4 (α-Syn^M5K,L8F,K10I,A17G,A19V^) or both EDC4 and membrane binding (α-Syn^Δ2–10^)^44^. Both variants were stabilized regardless of MpEDC4 expression, while wild-type α- Synuclein was degraded in the presence of MpEDC4 (Fig. 5D-E, Fig. S8C–D). These results confirm that α-Synuclein clearance depends on its interaction with EDC4.

We next asked whether human EDC4 could be engineered to acquire ATG8-binding capacity. To this end, we inserted the MpEDC4 AIM-containing region (residues D802–G906) into the intrinsically disordered region of HsEDC4 (between P677 and S707), generating a chimeric construct termed HsEDC4^AIM+^ (Fig. 5F). HEK293T cells were transfected with GFP-tagged versions of HsEDC4^WT^, HsEDC4^AIM+^, MpEDC4^WT^, or MpEDC4^AIM-^, along with mCherry-tagged GABARAP-L1. Co-IP assays showed that only MpEDC4^WT^ and HsEDC4^AIM+^ robustly interacted with GABARAP-L1 (Fig. 5C, Fig. S8B). This was confirmed in HEK293T cells stably expressing GFP-tagged constructs via lentiviral transduction (Fig. S8E). Confocal microscopy further revealed that HsEDC4^AIM+^, but not HsEDC4^WT^, formed puncta that co-localized with GABARAP-L1 (Fig. S8F).

We then assessed whether these constructs retain P-body localization. Unlike MpEDC4, which failed to interact with P-body markers HsDCP1A and HsXRN1, the HsEDC4^AIM+^ chimera bound both, recapitulating the behavior of wild-type HsEDC4 (Fig. 5G, Fig. S8G).

Finally, we tested whether the chimeric HsEDC4^AIM+^ could promote α-Synuclein degradation. Unexpectedly, it failed to do so and instead led to α-Synuclein accumulation (Fig. 5H, Fig. S8H). As controls, we transfected cells with MpEDC4^WT^ and MpEDC4^AIM-^. Only MpEDC4^WT^ promoted α-Synuclein clearance. These findings indicate that grafting an AIM is not sufficient to confer autophagy receptor function. We conclude that additional biochemical features of MpEDC4 - beyond ATG8 binding—are required to reprogram HsEDC4 into a functional autophagy receptor (Fig. 5I).

## Discussion

In this study, we identify MpEDC4 and MpDCP1 as a functional autophagy receptor pair that mediates selective degradation of P-bodies in the early land plant *Marchantia polymorpha*. Although both proteins are conserved components of the mRNA decapping machinery^1,45^, our findings reveal an additional, previously unrecognized role in promoting autophagic turnover of RNA granules. This function appears to be evolutionarily restricted, as the *Arabidopsis* and human orthologs of EDC4 and DCP1 do not associate with ATG8 or facilitate P-body degradation via autophagy. These results uncover an unexpected link between RNA metabolism and selective autophagy, revealing a lineage-specific functional innovation layered onto conserved RNA metabolism factors.

We demonstrate that MpEDC4 directly binds ATG8 through a canonical AIM motif within its intrinsically disordered region (IDR). While autophagic degradation of wild-type MpEDC4 was robust, we observed partial degradation of an AIM-deficient mutant (MpEDC4^Y845A^), suggesting the existence of additional or compensatory pathways. This mirrors observations in other selective autophagy systems—such as mitophagy and ER-phagy—where multiple receptors often act redundantly to ensure cargo recognition and clearance^46–48^.

Intriguingly, we show that MpDCP1 also functions as an ATG8 interactor, but through a non-canonical reverse AIM motif embedded in its own IDR. Mutation of two phenylalanine residues in this region abolished ATG8 binding, confirming its functional relevance. Dual mutation of AIM motifs in both MpEDC4 and MpDCP1 significantly impaired autophagic flux and compromised degradation of a third P-body component, DCP2. Together, these findings support a model in which the MpEDC4–MpDCP1 complex acts as a selective autophagy receptor module, coupling decapping machinery to autophagic clearance of RNA granules.

Beyond evolutionary insight, we showed functional activity across kingdoms. Expression of MpEDC4 in human cells promoted clearance of α-Synuclein, a protein implicated in Parkinson’s disease. Remarkably, engineering a chimeric HsEDC4 containing the MpEDC4 AIM motif (HsEDC4^AIM+^) restored ATG8 binding and P-body interactions, yet failed to trigger α-Synuclein degradation. These results suggest that ATG8 engagement alone is insufficient for receptor functionality, and that MpEDC4 contains additional molecular features— potentially involving structural elements, co-factors, or cargo recognition sequences—required for substrate degradation.

Collectively, our findings reveal MpEDC4 as a modular, dual-function protein that integrates mRNA decapping and selective autophagy. The ability of a liverwort-derived receptor to interface with human autophagy machinery highlights its potential as a synthetic degradation module. This work establishes a conceptual and technical foundation for engineering cross-kingdom autophagy receptors, with potential applications in targeted clearance of aggregation-prone proteins such as α-Synuclein. Future studies will be essential to define the biochemical determinants of MpEDC4 receptor competence and explore whether this approach can be extended to combat other disease-relevant substrates. Our study provides a framework for leveraging evolutionarily divergent autophagy components in synthetic biology and protein quality control.

## Acknowledgements

We thank BioOptics, Plant Sciences, Molecular Biology, Mass Spectrometry, Peptide Synthesis, Protein Technologies of The Vienna BioCenter Core Facilities (VBCF). We thank Sascha Martens, Stefan Ameres, Arturo Marí-Ordóñez, and members of the Dagdas lab for the fruitful discussions We thank Cécile Bousquet-Antonelli, Julia Bailey-Serres, Morten Petersen, Aurélien Boisson-Dernier and Takashi Ueda for sharing published material.

## Author contributions

AA designed the project, generated Arabidopsis and Marchantia lines, generated human HEK293 cell lines, performed flux experiments, microscopy, IP-MS experiments, phenotypic characterization experiments, wrote the manuscript. EE performed experiments with human cells. ASA and RKP analyzed data. NG generated Arabidopsis lines. EH, GEK, YFD secured funding, designed the project, supervised experiments, wrote and revised the manuscript.

## Data availability

The mass spectrometry proteomics data have been deposited to the ProteomeXchange Consortium via the PRIDE partner repository with the dataset identifier PXD064372. All the raw data are available via Zenodo (DOI: 10.5281/zenodo.15726897).

## Material and methods

### Yeast Two-Hybrid Screen

ULTImate Yeast Two-Hybrid Screen was performed by Hybrigenics Services (Paris, France). The *Marchantia polymorpha ATG8A* and *ATG8B* (https://marchantia.info/MpTak_v7.1/Mp1g21590/ and https://marchantia.info/MpTak_v7.1/Mp5g05930/) coding sequences were cloned into the pB66 as a C-terminal fusion with the DNA-binding domain of Gal4 (*N-Gal4-ATG8A-C* or *N- Gal4-ATG8B-C*). Those constructs were used as the baits for the screening of Marchantia cDNA library generated by random priming from *Marchantia polymorpha* gemmae thallus. A total of 62.7 million clones were screened for *ATG8A* and 43.1 million clones were screened for *ATG8B.* Positive clones were PCR-amplified and sequenced at 5’ and 3’ junctions. The resulting sequences were used to the corresponding interacting proteins in the Phytoxome13 JGI (https://phytozome-next.jgi.doe.gov/). A confidence score was shown for each interaction and ranked from A to F, where A is very high confidence in the interaction and F is proven technical artifacts (Supplementary Data 1).

### Plant material and growing conditions

All *Marchantia polymorpha* lines used in this study originate from the Takaragaike-1 (Tak-1, male) or Takaragaike-2 (Tak-2, female) ecotypes and are listed in the material section. Wild type plants were maintained and cultivated asexually through gemmae on half-strength ½ Gamborg B5 media (Gamborg B5 medium basal salt mixture supplemented with 0.5 g/L MES, pH 5.5) plates containing 1% plant agar under continuous white light with a light intensity of 50-60 µM/m²/sec at 22°C. Autophagy mutant plants were cultivated on the same media in a presence of 1% sucrose. For phenotyping and crossing of mature plants, 6-8 gemmae per line were grown on ½ Gamborg B5 plates in full white light for 10-12 days and then transferred to soil (Neuhaus Huminsubstrat from Klasmann-Deilmann GmbH and vermiculite in 2:1 proportion) inside plastic closed containers with air filters (produced by Sac02, Belgium) and grown under far-red-enriched irradiation. After approximately 4 weeks mature male and female reproductive organs developed. For fertilization, male reproductive organs were transferred to the petri dish and supplemented with 500 uL of sterile water to release spermatozoids, which were transferred to female reproductive organs. After approximately a month mature sporangia were visible and ready to be collected and stored at -80°C for longer storage.

All *Arabidopsis thaliana* lines used in this study originate from Columbia (Col-0) ecotype and are listed in the material section. Segregating *vcs-6* mutant seeds were obtained from SAIL (SAIL_831_D08, stock name CS837130). Primers used for genotyping are listed in the material section. Seeds were vapor-phase sterilized (90% sodium hypochlorite and 10% HCl) and sown on a water-saturated peat-based substrate (Klasmann Substrat 2, Klasmann-Deilmann GmbH). A *Bacillus thuringiensis* solution was used as a preventative control against fungus gnats (Gnatrol, Valent BioSciences, Libertyville, IL, USA). Lighting was provided by a custom-designed, multi-channel LED system developed in collaboration with RHENAC GreenTec AG (Germany), optimized for uniformity. The plants were cultured at 21°C in a 16- hour light/8-hour dark photoperiod under 165 µmol m-2 s-1 (PPFD) light intensity, using the following LED channels: 400 nm, white 4500K, 660 nm, and 730 nm.

### Carbon starvation treatments

12-days old *M.polymorpha* gemmae were transferred to the 1% agar plates supplemented with 0.5 Gamborg and MES, without sucrose, followed by transferring to either normal light irradiation or in the darkness for 2 days. *atg7-1* mutant plants were used as control

### Plant imaging

All plant pictures were acquired using a Canon EOS 80D DSLR equipped with either 60 mm fixed lenses or 18-135 mm lenses, using manual setting in CR2 format. The camera was mounted on a fixed stand and a dark cloth was used as background. Images were opened using Adobe Photoshop, where brightness and contrast was homogenized throughout all the images, and pictures were exported in tiff format to assemble panels.

### Human cell culture conditions

HEK Flp-In^TM^ T-REx^TM^ 293T cell line expressing doxycycline-inducible wild type α-synuclein was provided by Erinc Hallacli. The cells were maintained in Dulbecco’s modified Eagle’s Medium (DMEM) with 10% FBS, 1% L-Glutamine and 1% Penicillin/Streptomycin. Transfection was performed with jetOPTIMUS transfection reagent according to manufacturer’s instructions^49^. Cell lines were authenticated using STR profiling and repeatedly tested negative for mycoplasma contamination. Testing and authentication were performed using the in-house core facilities.

### Transfection of packaging cells

Plasmids used for transfection were purified using endotoxin-free in-house kit. Transfections were performed using 1100 ng of DNA in total, with 500 ng of plasmid of interest, 500 ng pCMVR8.74 (Addgene plasmid # 22036), and 100ng of pCMV-VSV-G (Addgene plasmid # 8454). Supplement free DMEM was used to mix DNA and Polyethylenimine (PEI) in a 1:3 ratio. HEK293T HiEX cell were used as packaging cells. 1×10^6^ cells were seeded in a 6-well plate a day before transfection and grown in a fully supplemented DMEM. The Plasmid mixture containing a transfection reagent was added dropwise onto the cells and they were incubated for 72h.

### Transduction of HEK293T cells

HEK Flp-In^TM^ T-REx^TM^ 293T expressing doxycycline-inducible wild type α-synuclein were seeded one day prior to transduction in fully supplemented media. 1×10^6^ cells were seeded in a 24-well plate. After 72h incubation, virus was collected from the supernatant with a syringe and sterile filtered. The sterile filtered virus was mixed with fully supplemented DMEM at the 1:10 ratio with 5ug/mL Polybrene. Cells were grown 2 rounds of 48h up to 10 cm dishes. After splitting cells in S2 conditions, cells were transferred into S1 conditions followed by FACS sorting gated for high and low expression. The high-expression population was used for further experiments.

### Cloning procedure

Coding sequences from genes of interest were amplified from cDNA with primers listed in the material section. The internal BsaI sites were mutated by site-directed-mutagenesis without affecting the amino acid sequence.

Constructs for *Marchantia polymorpha* were generated using either the OpenPlant toolkit^50^ or GreenGate^51^ cloning systems. Transgenic lines were generated by Agrobacterium-mediated transformation of regenerated thalli as previously described^52^. Briefly, ∼4cm2 M. polymorpha thalli extracted from 14-d-old plants were co-incubated with Agrobacterium containing the desired constructs for 2 days in transformation media (1.5 g/L Gamborg B5 medium, 2.5 mM MES monohydrate, 20 g/L D-Sucrose, 1g/L Casein hydrolysate, 2 mM L-Glutamine) supplemented with 100 µM Acetosyringone. Transformants were selected based on antibiotic resistance on half-strength Gamborg’s B5 medium containing 1% agar without sucrose supplemented with Ticarcillin.

Stable transgenic *Arabidopsis thaliana* lines were generated by Agrobacterium-mediated transformation using the floral dip method^53^.

Constructs for *Escherichia coli* transformation were assembled with the GreenGate^51^ cloning system. All plasmids used in this study, along with the corresponding primers used for their generation, are listed in in the material section.

### Autophagy flux assay in *Marchantia polymorpha*

For Torin-1 autophagic flux assay, 6 days old *Marchantia polymorpha* gemmaes were incubated in either DMSO or 12 μM Torin-1-containing media (0.5 Gamborg, MES, 1% sucrose) for 4 h, after which plants were either harvested or recovered for additional 20 h with either fresh media or media containing 10 μM E64d and 10 μM pepstatin A before harversing. For high salt flux assay, 6-days old *Marchantia polymorpha* gemmaes were incubated in either fresh media or 150 mM sodium-chloride (NaCl) containing media (0.5 Gamborg, MES, 1% sucrose) for 4 h, after which plants were recovered for additional 20 h with either fresh media or media containing 10 μM E64d and 10 μM pepstatin A before harversing.

For affinity purification, *Marchantia polymorpha* gemmaes were cultured in liquid half-strength Gamborg’s B5 medium with 1% sucrose for 10 days without shaking.

### Autophagy flux assay in *Arabidopsis thaliana*

30-40 seedlings for western blot were grown on the plates (half-strength MS media (Murashige and Skoog salt + Gamborg B5 vitamin mixture) with 1% sucrose, 0.5 g/L MES and 1% plant agar) 6 days under continuous light. One day prior the treatment, seedlings were transferred to the liquid media (half-strength MS media (Murashige and Skoog salt + Gamborg B5 vitamin mixture) with 1% sucrose, 0.5 g/L MES). On the 7^th^ day, seedlings were incubated in either DMSO or 3 μM Torin-1-containing media, after which plants were either harvested or recovered for additional 4h and 20 h with either fresh media or media containing 1 μM concanamycin A before harvesting.

### Western blotting

Plant material was harvested following the respective treatments and flash-frozen in liquid nitrogen. Frozen tissue was ground to a fine powder in a 1.5 Safe-Lock Eppendorf tubes with glass beads (Carl Roth; 1,7-2,1 mm) using a Silamat S6. Total protein extraction was achieved by adding 350 µl of 2X Laemmli buffer to the ground tissue and mixing in the Silamat S6 for 20 seconds until the ground tissue and the buffer were homogenized. Samples were denatured at 70°C for 10 min and centrifuged for 5 minutes at maximum speed in a microcentrifuge. Protein quantification was performed with Bradford assay (Sigma-Aldrich) following the manufacturer’s instructions or with the Amidoblack method. Briefly, 10 µl of the protein sample was diluted in 190 µl of deonized water and mixed thoroughly with 1 ml of normalized Amidoblack staining solution (90% methanol, 10% acetic acid, 0.05% Naphtol Blue Black). Samples were centrifuged at maximum speed for 10 minutes. Supernatant was discarded and the resulting pellets were washed with 1 ml of washing solution (90% ethanol and 10% acetic acid), centrifuged at maximum speed for 10 minutes and dissolved in 1 ml 0.2 N NaOH. The corresponding optical density (OD) at 630 nm was measured in a plate reader (Synergy HTX Multi-Mode Microplate Reader; BioTek) using NaOH solution as blank. Protein concentration was calculated using the formula C = (OD-b)/10a, where a and b are calibrated by Bovine Serum Albumin (BSA) standard curve of the staining solution. For Western blotting, the indicated total protein amount was loaded on SDS-PAGE gels (4-20% Mini-PROTEAN TGX precast gel; Bio-Rad) and blotted on nitrocellulose membrane (Bio-Rad) using the semi-dry Trans-Blot Turbo Transfer System (Bio-Rad). Membranes were blocked in TBST (10 mM Tris-HCl pH 7.5, 150 mM NaCl, 0.1% Tween 20) + 5% skimmed milk at room temperature for 1 hour. This was followed by an overnight incubation at 4°C with the primary antibody diluted in blocking buffer. After several washes with TBST, membranes were incubated for 45 min with the respective secondary antibody diluted in TBST + 5% skimmed milk and subsequently washed before imaging. All primary and secondary antibodies are provided in the materials section. The immune-reaction was developed using either Pierce™ ECL Western Blotting Substrate (ThermoFisher) or SuperSignal™ West Pico PLUS Chemiluminescent Substrate (ThermoFisher) and detected with either ChemiDoc Touch Imaging System (Bio-Rad) or iBright Imaging System (Invitrogen). Equal total protein loading on the membrane was corroborated by Amidoblack staining.

### Western blot image quantification

Protein bands intensities were quantified via Fiji software (version: 1.52 and later 2.14.0/1.54f). Equal rectangles were drawn around the total protein gel lane and the band of interest. The lane profile was obtained by subtracting the mean intensity of the background. The adjusted volume of the peak in the profile was taken as a measure of the band intensity. The protein band of interest was normalized for the total protein level of the whole lane. Average relative intensities and a standard error of at least three independent experiments were calculated.

### Affinity purification coupled to mass spectrometry (AP-MS) of *Marchantia polymorpha* samples

About 1-2 grams of plant material were harvested and homogenized using liquid nitrogen and dissolved in 1.5 volumes of lysis buffer (45 mM Tris-HCl pH 7.5, 150 mM NaCl, 10% glycerol, 1 mM EDTA, 0.15% Nonidet P-40, 1% PVPP, 1 mM DTT, Protease Inhibitor Cocktail tablet, RNasin Ribonuclease Inhibitor). Plant lysates were cleared by centrifugation at 16,000 x g for 15 at 4°C twice. Protein concentration was measured using Bradford protein assay (Sigma). Beads were equilibrated with pre-chilled IP buffer (45 mM Tris-HCl pH 7.5, 150 mM NaCl, 10% glycerol, 1 mM EDTA, 0.15% Nonidet P-40, 1 mM DTT). For peptide competition, normalized plant extracts were spiked with AIM wt or AIM mut peptides dissolved in IP buffer at a final concentration of 100 μM and incubated with 30 μl GFP-Trap or RFP-trap Magnetic Agarose beads (ChromoTek) for 1 hour 30 min at 4°C. Pellets were washed for 5 times with detergent-free IP buffer (45 mM Tris-HCl pH 7.5, 150 mM NaCl, 10% glycerol, 1 mM EDTA) and submitted for mass spectrometry. To test whether IP worked, a small fraction of beads (10%) was boiled for 5 min at 95°C prior to immunoblotting with the respective antibodies. Mass spectrometry sample preparation and measurement were performed as previously described^30,31^.

### *In vivo* co-immunoprecipitation for human cells

About 2×10^6^ HEK293T cells were seeded in a 6-well plate in a fully supplemented DMEM. After 24h incubation, cells were transfected with jetOPTIMUS transfection reagent according to manufacturer’s instructions^49^. After 48h after transfection, cells were harvested using approximately 200 μl of lysis buffer (45 mM Tris-HCl pH 7.5, 150 mM NaCl, 10% glycerol, 1 mM EDTA, 0.5% Nonidet P-40, 1% PVPP, 1 mM DTT, Protease Inhibitor Cocktail tablet, RNasin Ribonuclease Inhibitor). Lysates were cleared by centrifugation at 16,000 x g for 15 at 4°C. Protein concentration was measured using Bradford protein assay (Sigma). Beads were equilibrated with pre-chilled IP buffer (45 mM Tris-HCl pH 7.5, 150 mM NaCl, 10% glycerol, 1 mM EDTA, 0.2% Nonidet P-40, 1 mM DTT). Final lysates were incubated with 30 μl GFP-Trap or RFP-trap Magnetic Agarose beads (ChromoTek) for 1 hour 30 min at 4°C. Pellets were washed for 5 times with IP buffer following by boiling for 5 min at 95°C prior to immunoblotting with the respective antibodies.

### *In vitro* pulldowns

For pulldown experiments, 10 µl of glutathione magnetic agarose beads (Pierce Glutathione Magnetic Agarose Beads, Thermo Scientific) were equilibrated by washing them twice with wash buffer (100 mM NaPi pH 7.2, 300 mM NaCl, 1 mM DTT, 0.01% (v/v) Nonidet P-40). Normalized E. coli clarified lysates were mixed, added to the washed beads, and incubated on an end-over-end rotator for 1 hour at 4°C. Beads were washed five times in 1 ml wash buffer. Bound proteins were eluted by adding 35 µl Laemmli buffer. Samples were analyzed by western blotting.

### Sample preparation for confocal microscopy

For *Marchantia polymorpha* confocal microscopy, 2-d-old thalli were placed on a microscope slide with water and covered with a coverslip. The meristematic region was used for image acquisition.

Transfected human cells were grown on coverslips and fixed utilizing 0.4% Paraformaldehyde solution in PBS for 15 min, followed by 3 washed with cold PBS. Fixed cells were mounted in VectaShield mounting medium without DAPI and sealed using clear nail polish.

### Confocal microscopy

Samples were imaged at an upright ZEISS LSM800 or LSM 780 confocal microscope (Zeiss) with an Apochromat 40x or 63x objective lens at 1x magnification. mKalama1 fluorescence was excited at 405 nm and detected at 420 and 470 nm. YFP or GFP fluorescence was excited at 488 nm and detected between 510 and 546 nm. mScarlet or mCherry fluorescence was excited at 561 nm and detected between 571 and 617 nm. For each experiment, all replicate images were acquired using identical parameters. Confocal images were processed with Fiji (version 1.52, Fiji) and exported as .tiff for panel assembly

### Image processing and quantification

For co-localization experiments, single snaps were used for quantification. For co-localization experiments, Mander’s co-localization analyses were performed via Fiji software (version: 1.52 and later 2.14.0/1.54f). M1 and M2 Mander’s coefficient values were calculated via the JACoP plugin. Thresholding for each Channel were adjusted for each snap image according to the puncta signal in original confocal images via JACoP plugin settings. Values near one represent almost perfect correlation, whereas values near 0 reflect no correlation

### Phylogenetic analyses

To build phylogeny of EDC4 and DCP1, protein sequences of EDC4 and DCP1 orthologs were collected from UniProt (UniProtKB). Sequences were aligned using Clustal Omega server^54^. Maximum-likelihood phylogenetic trees were constructed using MEGA software^55^, both with and without 500 bootstrap replicates.

### Statistical analysis

Statistical analyses were performed with GraphPad Prism eight software. For all the quantifications described above, statistical analysis was performed. Statistical significance of differences between two experimental groups was assessed wherever applicable by either a two-tailed Student’s t-test if the variances were not significantly different according to the F test or using a non-parametric test (Mann-Whitney or Kruskal-Wallis with Dunn’s post-hoc test for multiple comparisons) if the variances were significantly different (p<0.05). Differences between two data sets were considered significant at p<0.05 (*); p<0.01 (**); p<0.001 (***); p<0.0001 (****); n.s., not significant.

### Experimental Model/Cell lines

| Organism | Background |
| --- | --- |
| *Arabidopsis thaliana* | Col-0 |
| *Marchantia polymorpha* | Takaragaike-1 (Tak-1) |
| *Marchantia polymorpha* | Takaragaike-2 (Tak-2) |
| *Homo Sapiens* | HEK293T cell line |

### Plasmids

| Organism | Name | Reference |
| --- | --- | --- |
| *E. coli* | E. coli destination vector (pET) | 56 |
| *E. coli* | pET-GST | 56 |
| *E. coli* | pET-MBP | 56 |
| *E. coli* | pET-GST-AtATG8A | 56 |
| *E. coli* | pET-GST-AtATG8E | 56 |
| *E. coli* | pET-GST-MpATG8A | 56 |
| *E. coli* | pET-GST-MpATG8B | 56 |
| *E. coli* | pET-GST-MpATG8A<sup>ADS(Y52A,L53A)</sup> | 56 |
| *E. coli* | pET-GST-MpATG8B<sup>ADS(K51A)</sup> | 56 |
| *E. coli* | pET-GST-AtVCS | This study |
| *E. coli* | pET-GST-AtDCP1 | This study |
| *E. coli* | pET-GST-MpEDC4 linker | This study |
| *E. coli* | pET-GST-MpEDC4 linker<sup>F718A</sup> | This study |
| *E. coli* | pET-GST-MpEDC4 linker<sup>Y725A</sup> | This study |
| *E. coli* | pET-GST-MpEDC4 linker<sup>Y845A</sup> | This study |
| *E. coli* | pET-GST-MpEDC4 linker<sup>Y951A</sup> | This study |
| *E. coli* | pET-GST-MpDCP1 | This study |
| *E. coli* | pET-GST-MpDCP1<sup>F73A</sup> | This study |
| *E. coli* | pET-GST-MpDCP1<sup>F130A</sup> | This study |
| *E. coli* | pET-GST-MpDCP1<sup>F137A</sup> | This study |
| *E. coli* | pET-GST-MpDCP1<sup>F154A</sup> | This study |
| *E. coli* | pET-GST-MpDCP1<sup>F191A,F192A</sup> | This study |
| *E. coli* | pET-GST-MpDCP1<sup>F263A</sup> | This study |
| *A. thaliana* | pGGSun-Ub10::GFP-AtVCS | This study |
| *A. thaliana* | pGGSun-Ub10::GFP-MpEDC4 | This study |
| *A. thaliana* | pGGSun-Ub10::GFP-MpEDC4<sup>Y845A</sup> | This study |
| *H. sapiens* | pLX317 GFP | Addgene<br>(#193681) |
| *H. sapiens* | pLX317 GFP-HsEDC4 | This study |
| *H. sapiens* | pLX317 GFP-HsEDC4 aimPLUS | This study |
| *H. sapiens* | pLX317 GFP-MpEDC4 | This study |
| *H. sapiens* | pLX317 GFP-MpEDC4<sup>Y845A</sup> | This study |
| *H. sapiens* | pmCherry-Gabarap-L1-mCherry | 56 |
| *H. sapiens* | pEGFP(N2) | 56 |
| *H. sapiens* | pEGFP(N2)-HsEDC4 | This study |
| *H. sapiens* | pEGFP(N2)-HsEDC4 aimPLUS | This study |
| *H. sapiens* | pEGFP(C1)-MpEDC4 | This study |
| *H. sapiens* | pEGFP(C1)-MpEDC4<sup>Y845A</sup> | This study |
| *H. sapiens* | pcDNA5/FRT-TO/GW/humanKozak_SNCA_STOP_WT_3xFLAG-V5 | Erinc Hallacli |
| *H. sapiens* | pcDNA_FRT_TO_SNCA(D2-10)-STOP | Erinc Hallacli |
| *H. sapiens* | pcDNA_FRT_TO_SNCA_M5K_L8F_K10I_A17G_A19V_STOP | Erinc Hallacli |
| *M. polymorpha* | pCsA (L2_Hyg_lacZgRNA-Cas9) | 50 |
| *M. polymorpha* | pCsA (L2_Cs_lacZgRNA-Cas9) | 50 |
| *M. polymorpha* | pMpGWB303-pEF1α::GFP | 31 |
| *M. polymorpha* | pMpGWB303-pEF1α::GFP-MpATG8A | 31 |
| *M. polymorpha* | pMpGWB303-pEF1α::GFP-MpATG8B | 31 |
| *M. polymorpha* | pMpGWB303-pEF1α::GFP-MpATG8B<sup>ADS(K51A)</sup> | 31 |
| *M. polymorpha* | pGGSun-p35S::GFP-MpATG8A | This study |
| *M. polymorpha* | pGGSun-p35S::GFP-MpATG8B | This study |
| *M. polymorpha* | pMpGWB303-pEF1α::mScarlet | 31 |
| *M. polymorpha* | pGGSun-pEF1α::mScarlet-MpATG8B | This study |
| *M. polymorpha* | pGGSun-pEF1α::mScarlet-MpNBR1 (Mp3g25290) | This study |
| *M. polymorpha* | pGGSun-pEF1α::mScarlet-MpEDC4 | This study |
| *M. polymorpha* | pGGSun-pEF1α::mScarlet-MpEDC4<sup>Y845A</sup> | This study |
| *M. polymorpha* | pGGSun-pEF1α::GFP-MpDCP1 | This study |
| *M. polymorpha* | pGGSun-pEF1α::GFP-MpDCP2 | This study |
| *M. polymorpha* | pGGSun-pEF1α::GFP-MpATG11 | This study |
| *M. polymorpha* | pGGSun-pEDC4::GFP-MpEDC4 | This study |
| *M. polymorpha* | pGGSun-pEF1::MpDCP1-mKalama1 | This study |
| *M. polymorpha* | pGGSun-pEF1α::mScarlet-MpEDC4-P2A-MpDCP1-mKalama1 | This study |
| *M. polymorpha* | pGGSun-pEF1α::mScarlet-MpEDC4<sup>Y845A</sup>-P2A-MpDCP1<sup>F191A,F192A</sup>-mKalama1 | This study |

### Oligonucleotides

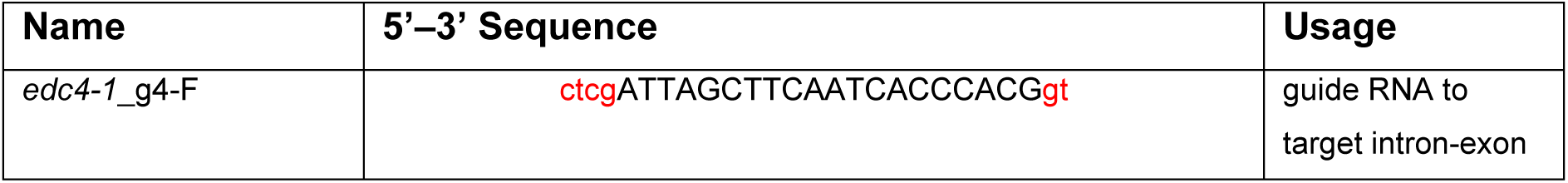
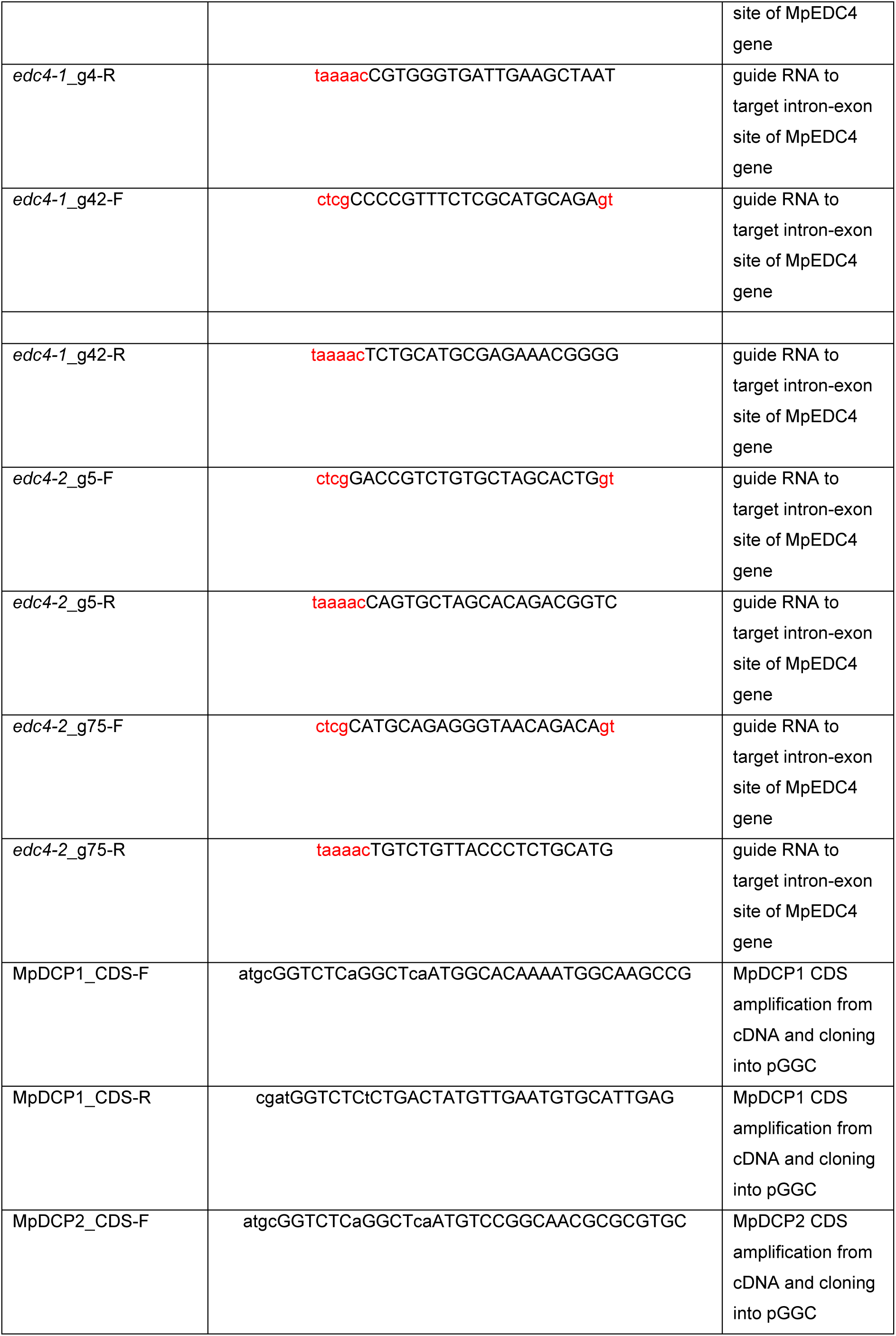
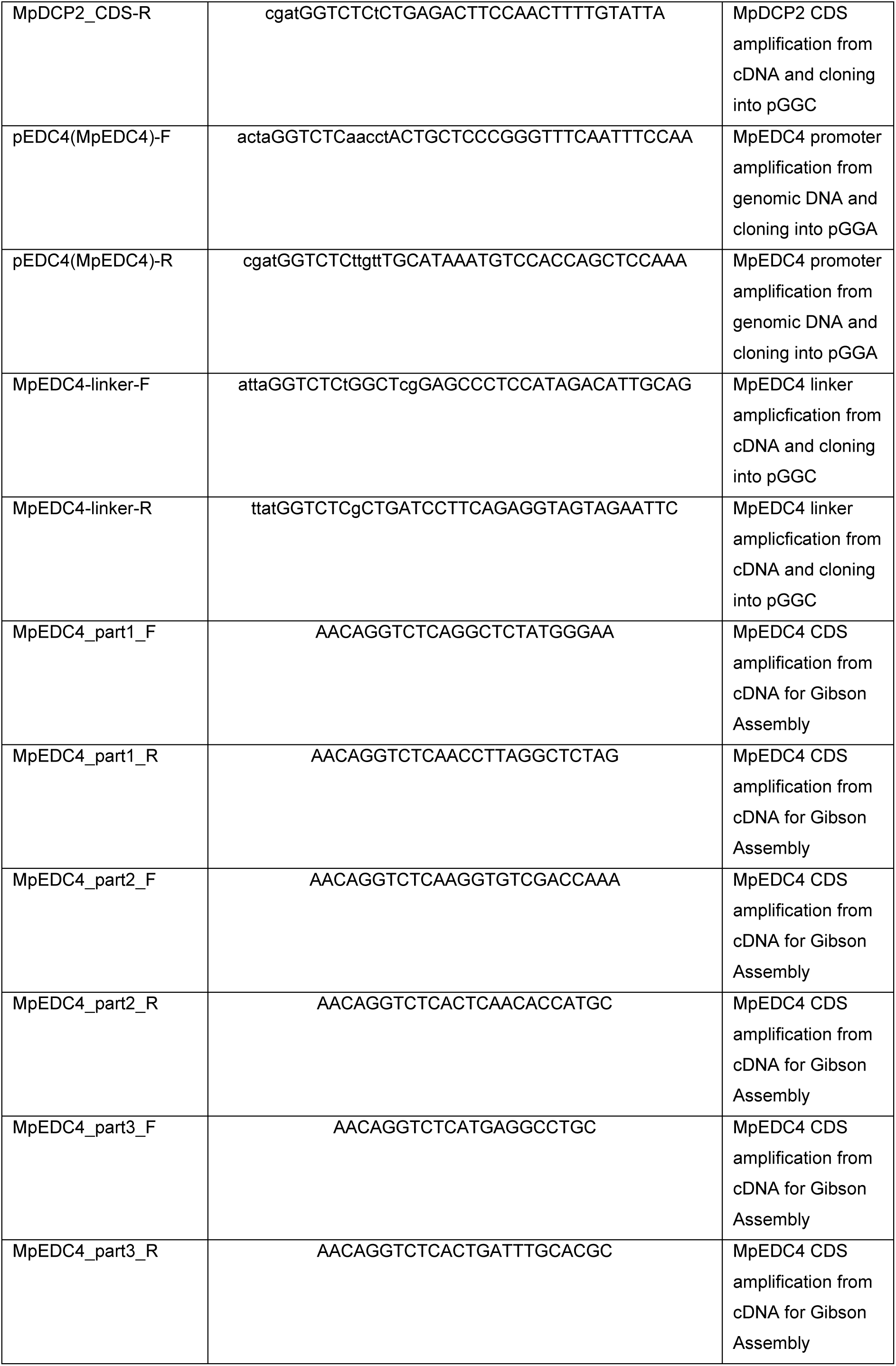
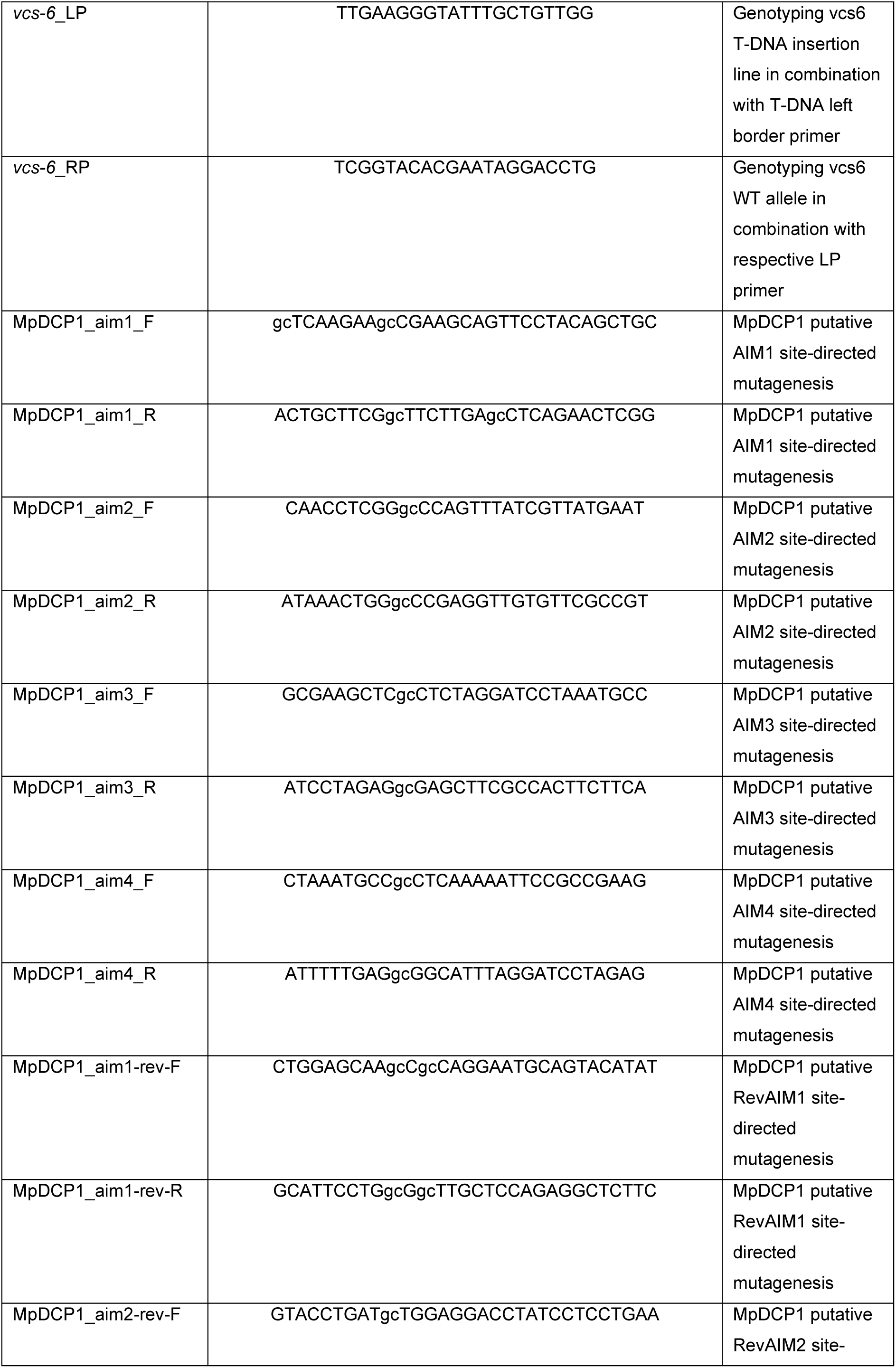
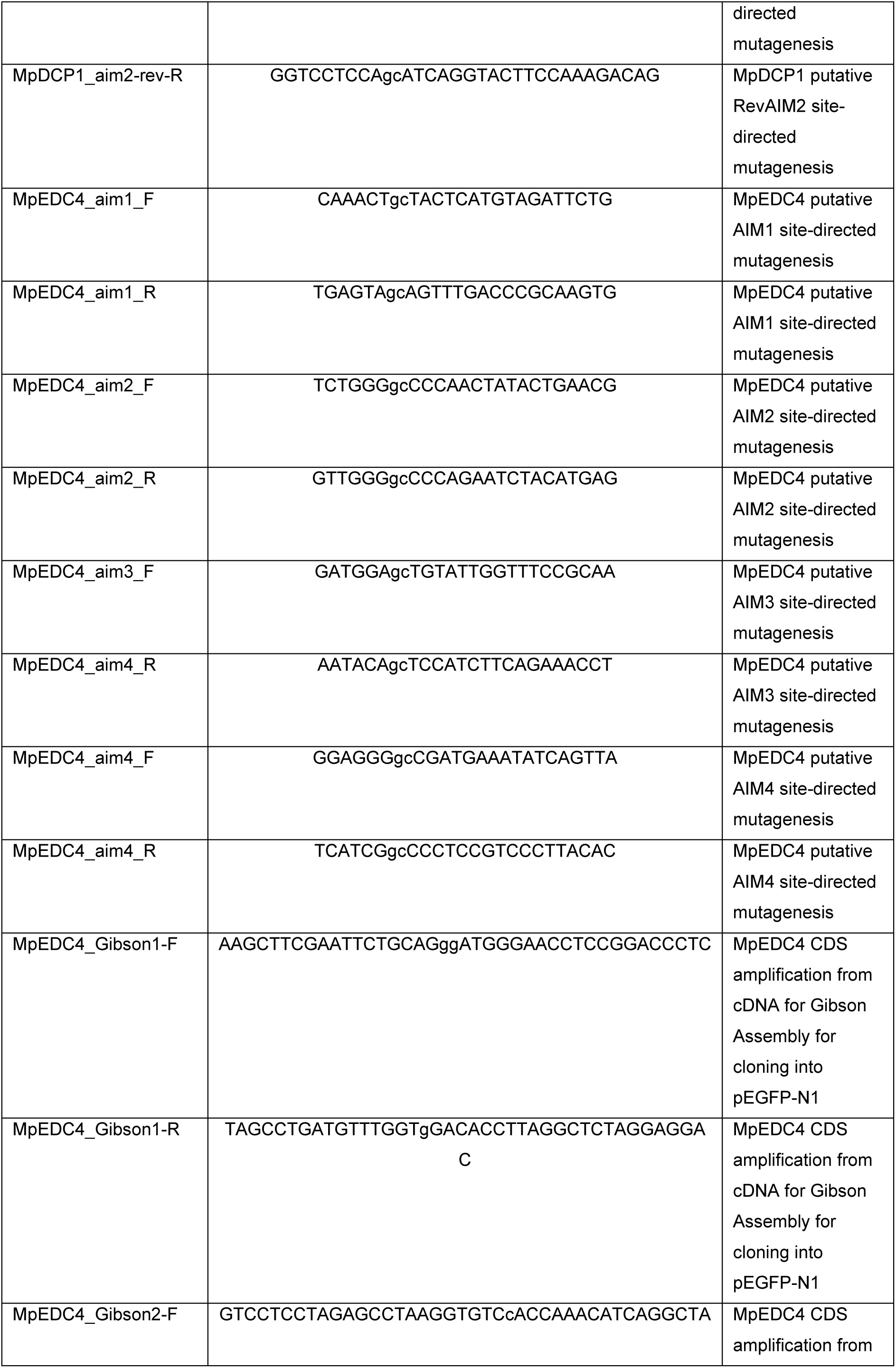
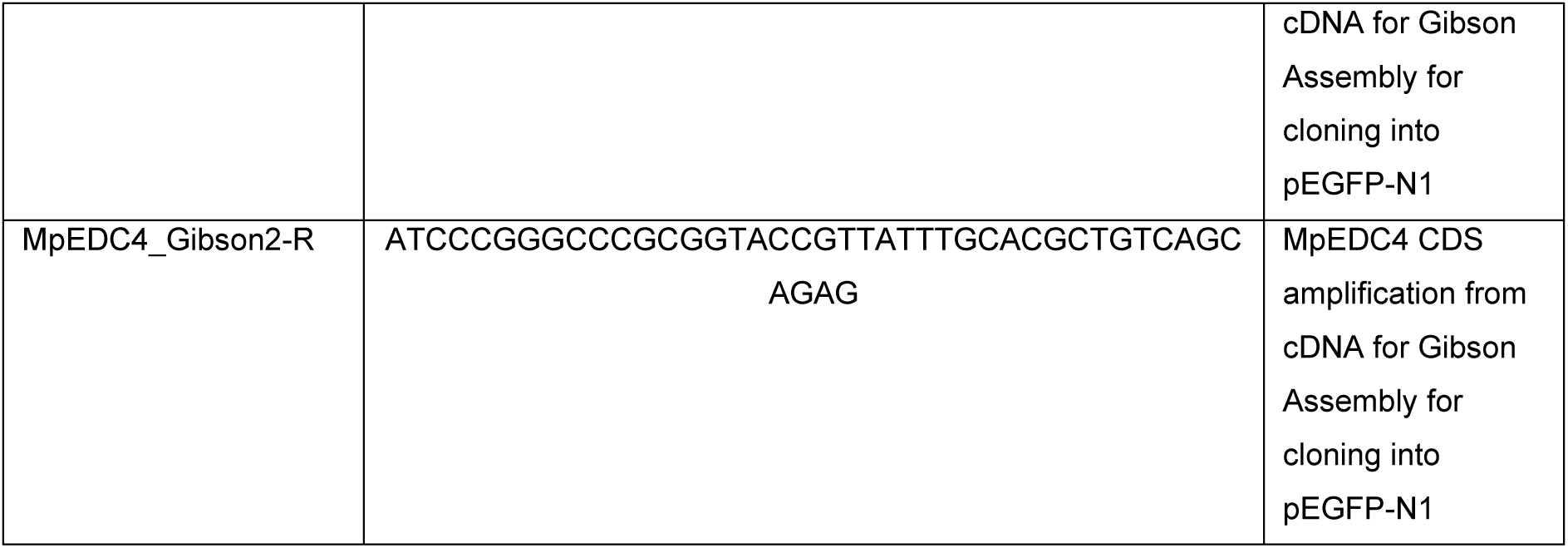

### Mutants

| Organism | Genotype | Gene ID | Preparation/T-DNA line | Supplier | Reference |
| --- | --- | --- | --- | --- | --- |
| *M. polymorpha* | *atg7-1* | Mp2g07850 | CRISPR |  | <sup>57</sup> |
| *A. thaliana* | *atg5-1* | AT5G17290 | SAIL_129B07 | NASC (N39993) | <sup>58</sup> |
| *A. thaliana* | *vcs-6* | AT3G13300 | SAIL_831_D08 | NASC (N837130) | <sup>59</sup> |

### Plant Lines

| Name | Accession number | Reference | Plant species |
| --- | --- | --- | --- |
| pEF1::GFP |  | <sup>31</sup> | *M. polymorpha* |
| pEF1::mScarlet |  | <sup>31</sup> | *M. polymorpha* |
| pEF1::GFP-MpATG8A | Mp1g21590 | <sup>31</sup> | *M. polymorpha* |
| pEF1::GFP-MpATG8B | Mp5g05930 | <sup>31</sup> | *M. polymorpha* |
| pEF1::GFP-MpATG8B<sup>ADS</sup> | Mp5g05930 | <sup>31</sup> | *M. polymorpha* |
| pEF1::mScarlet-MpATG8B | Mp5g05930 | This study | *M. polymorpha* |
| pEF1::mScarlet-MpEDC4 | Mp3g11710 | This study | *M. polymorpha* |
| pEF1::mScarlet-MpEDC4 in pEF1::GFP | Mp3g11710 | This study | *M. polymorpha* |
| pEF1::mScarlet-MpEDC4 in pEF1::GFP-MpATG8A | Mp3g11710 (EDC4)<br>Mp1g21590 (ATG8A) | This study | *M. polymorpha* |
| pEF1::mScarlet-MpEDC4 in pEF1::GFP-MpATG8B | Mp3g11710 (EDC4)<br>Mp5g05930 (ATG8B) | This study | *M. polymorpha* |
| pEF1::mScarlet-MpEDC4 in<br>pEF1::GFP-MpATG8B<sup>ADS</sup> | Mp3g11710 (EDC4)<br>Mp5g05930 (ATG8B) | This study | *M. polymorpha* |
| pEDC4<sub>min</sub>::GFP-EDC4 in<br>pEF1::mScarlet-ATG8B | Mp3g11710 (EDC4)<br>Mp5g05930 (ATG8B) | This study | *M. polymorpha* |
| pEF1::mScarlet-MpEDC4 in<br>pEF1::GFP-MpATG11 | Mp3g11710 (EDC4)<br>Mp4g18080 (ATG11) | This study | *M. polymorpha* |
| pEF1::mScarlet-EDC4 in *atg7-1* | Mp3g11710 (EDC4)<br>Mp2g07850 (ATG7) | This study | *M. polymorpha* |
| pEF1::mScarlet-EDC4 in *edc4-1* | Mp3g11710 (EDC4) | This study | *M. polymorpha* |
| pEF1::mScarlet-EDC4 in *edc4-2* | Mp3g11710 (EDC4) | This study | *M. polymorpha* |
| pEF1::mScarlet-MpEDC4 in pEF1::<br>GFP-DCP1 | Mp3g11710 (EDC4)<br>Mp1g25660 (DCP1) | This study | *M. polymorpha* |
| pEF1::mScarlet-MpEDC4 in pEF1::<br>GFP-DCP2 | Mp3g11710 (EDC4)<br>Mp8g16420 (DCP2) | This study | *M. polymorpha* |
| pEF1::GFP-DCP1 | Mp1g25660 (DCP1) | This study | *M. polymorpha* |
| pEF1::GFP-DCP1 in<br>pEF1::mScarlet-ATG8B | Mp1g25660 (DCP1)<br>Mp5g05930 (ATG8B) | This study | *M. polymorpha* |
| pEF1::GFP-DCP1 in *atg7-1* | Mp1g25660 (DCP1)<br>Mp2g07850 (ATG7) | This study | *M. polymorpha* |
| pEF1::GFP-DCP2 | Mp8g16420 (DCP2) | This study | *M. polymorpha* |
| pEF1::GFP-DCP2 in *atg7-1* | Mp8g16420 (DCP2)<br>Mp2g07850 (ATG7) | This study | *M. polymorpha* |
| pEF1::mScarlet-MpEDC4<sup>Y845A</sup> | Mp3g11710 (EDC4) | This study | *M. polymorpha* |
| pEF1::mScarlet-MpEDC4<sup>Y845A</sup> in<br>pEF1::GFP | Mp3g11710 (EDC4) | This study | *M. polymorpha* |
| pEF1::mScarlet-MpEDC4<sup>Y845A</sup> in<br>pEF1::GFP-MpATG8A | Mp3g11710 (EDC4)<br>Mp1g21590 (ATG8A) | This study | *M. polymorpha* |
| pEF1::mScarlet-MpEDC4<sup>Y845A</sup> in<br>pEF1::GFP-MpATG8B | Mp3g11710 (EDC4)<br>Mp5g05930 (ATG8B) | This study | *M. polymorpha* |
| pEF1::mScarlet-EDC4<sup>Y845A</sup> in<br>*edc4-1* | Mp3g11710 (EDC4) | This study | *M. polymorpha* |
| pEF1::mScarlet-EDC4<sup>Y845A</sup> in<br>*edc4-2* | Mp3g11710 (EDC4) | This study | *M. polymorpha* |
| pEF1::DCP1-mKalama1-3xflag in<br>pEF1::mScarlet-MpEDC4 +<br>pEF1::GFP-MpATG8A | Mp3g11710 (EDC4)<br>Mp1g25660 (DCP1) | This study | *M. polymorpha* |
| pEF1::GFP-NBR1 in<br>pEF1::mScarlet-MpEDC4 + DCP1-mKalama1-3xflag | Mp3g11710 (EDC4)<br>Mp1g25660 (DCP1)<br>Mp3g25290 (NBR1) | This study | *M. polymorpha* |
| pEF1::DCP1-mKalama1-3xflag-P2A-mScarlet-MpEDC4 | Mp3g11710 (EDC4)<br>Mp1g25660 (DCP1) | This study | *M. polymorpha* |
| pEF1::DCP1<sup>F191A,F192A</sup>-mKalama1-3xflag-P2A-mScarlet-MpEDC4<sup>Y845A</sup> | Mp3g11710 (EDC4)<br>Mp1g25660 (DCP1) | This study | *M. polymorpha* |
| pEF1::DCP1-mKalama1-3xflag-P2A-mScarlet-MpEDC4 in<br>pEF1::GFP-ATG8A | Mp3g11710 (EDC4)<br>Mp1g21590 (ATG8A)<br>Mp1g25660 (DCP1) | This study | *M. polymorpha* |
| pEF1::DCP1<sup>F191A,F192A</sup>-mKalama1-3xflag-P2A-mScarlet-MpEDC4<sup>Y845A</sup><br>in pEF1::GFP-ATG8A | Mp3g11710 (EDC4)<br>Mp1g21590 (ATG8A)<br>Mp1g25660 (DCP1) | This study | *M. polymorpha* |
| pEF1::DCP1-mKalama1-3xflag-P2A-mScarlet-MpEDC4 in<br>pEF1::GFP-DCP2 | Mp3g11710 (EDC4)<br>Mp1g25660 (DCP1)<br>Mp8g16420 (DCP2) | This study | *M. polymorpha* |
| pEF1::DCP1<sup>F191A,F192A</sup>-<br>mKalama1-3xflag-P2A-mScarlet-MpEDC4<sup>Y845A</sup> in<br>pEF1::GFP-DCP2 | Mp3g11710 (EDC4)<br>Mp1g25660 (DCP1)<br>Mp8g16420 (DCP2) | This study | *M. polymorpha* |
| pUBQ::GFP-AtVCS in *vcs-6* | AT3G13300 (VCS) | This study | *A. thaliana* |
| pUBQ::GFP-MpEDC4 wt in *vcs-6* | Mp3g11710 (EDC4)<br>AT3G13300 (VCS) | This study | *A. thaliana* |
| pUBQ::GFP-MpEDC4<sup>Y845A</sup> in *vcs-6* | Mp3g11710 (EDC4)<br>AT3G13300 (VCS) | This study | *A. thaliana* |
| pDCP1::YFP-DCP1 | AT1G08370 (DCP1) | <sup>60</sup> | *A. thaliana* |
| pDCP1::YFP-DCP1 x *atg5-1* | AT1G08370 (DCP1)<br>AT5G17290 (ATG5) | This study | *A. thaliana* |
| pDCP1::YFP-DCP1 x<br>pUBQ::mCherry-AtATG8E | AT1G08370 (DCP1)<br>AT2G45170 (ATG8E) | This study | *A. thaliana* |

### Human cell lines

| Name | Accession number | Reference | Cell line |
| --- | --- | --- | --- |
| GFP |  | This study | HEK293T |
| GFP-HsEDC4 *wt* | Q6P2E9 | This study | HEK293T |
| GFP-HsEDC4 *aimPLUS* | Q6P2E9 | This study | HEK293T |
| GFP-MpEDC4 *wt* | Mp3g11710 | This study | HEK293T |
| GFP-MpEDC4<sup>Y845A</sup> | Mp3g11710 | This study | HEK293T |

### Bacterial Strains

| Organism | Strain | Supplier |
| --- | --- | --- |
| *E. coli* | DH5α | In-house facility |
| *E. coli* | Rosetta2 (DE3) pLysS | In-house facility |
| *A. tumefaciens* | GV3101 (pSoup) | In-house facility |

### Peptides

| Name | Source | Sequence |
| --- | --- | --- |
| AIM *wt* | Synthesized *in house* | EPLDFDWEIVLEEEM |
| AIM *mut* | Synthesized *in house* | EPLDFDAEIALEEEM |

### Antibodies

**Antibodies:**
| Antibody | Host | Supplier | Cat. # | Working concentration |
| --- | --- | --- | --- | --- |
| Anti-Rabbit IgG HRP-Conjugate | Goat | Bio-Rad | 1706515 | 1:5000 |
| Anti-Mouse IgG-HRP Conjugate | Goat | Bio-Rad | 1706516 | 1:5000 |
| Anti-GFP | Mouse | Roche | 11814460001 | 1:2500 |
| Anti-GFP | Rabbit | Invitrogen | A11122 | 1:2500 |
| Anti-mCherry | Rabbit | Proteintech | 26765-1-AP | 1:3000 |
| Anti-GST HRP-Conjugate | Goat | GE Healthcare | RPN1236 | 1:5000 |
| Anti-MBP | Mouse | Sigma Aldrich | M1321 | 1:5000 |
| Anti-NBR1 | Rabbit | Agrisera | AS14 2805 | 1:5000 |
| Anti-α-Synuclein | Rabbit | Cell Signaling Technology | E4U2F | 1:1000 |
| Anti-DCP1A | Rabbit | Proteintech | 22373-1-AP | 1:2000 |
| Anti-XRN1 | Mouse | Santa Cruz | sc-165985 | 1:2000 |

### Inhibitors and drugs

| Name | Supplier | Cat. # |
| --- | --- | --- |
| Concanamycin A (ConA) | Santa Cruz Biotechnology | sc-202111A |
| E64d | Santa Cruz Biotechnology | sc-201280 |
| Pepstatin A | Santa Cruz Biotechnology | sc-45036 |
| Torin1 (Torin) | Santa Cruz Biotechnology | sc-396760 |
| Doxycycline | Sigma-Aldrich | 24390-14 |

### Affinity matrices for purification and immuno-precipitation

| Name | Supplier | Cat. # |
| --- | --- | --- |
| GFP-Trap Magnetic Agarose | Chromotek | Gtma |
| RFP-Trap Magnetic Agarose | Chromotek | Rtma |
| Pierce™ Glutathione Magnetic Agarose Beads | Thermo Scientific™ | 78601 |

---

**Figure S1.**
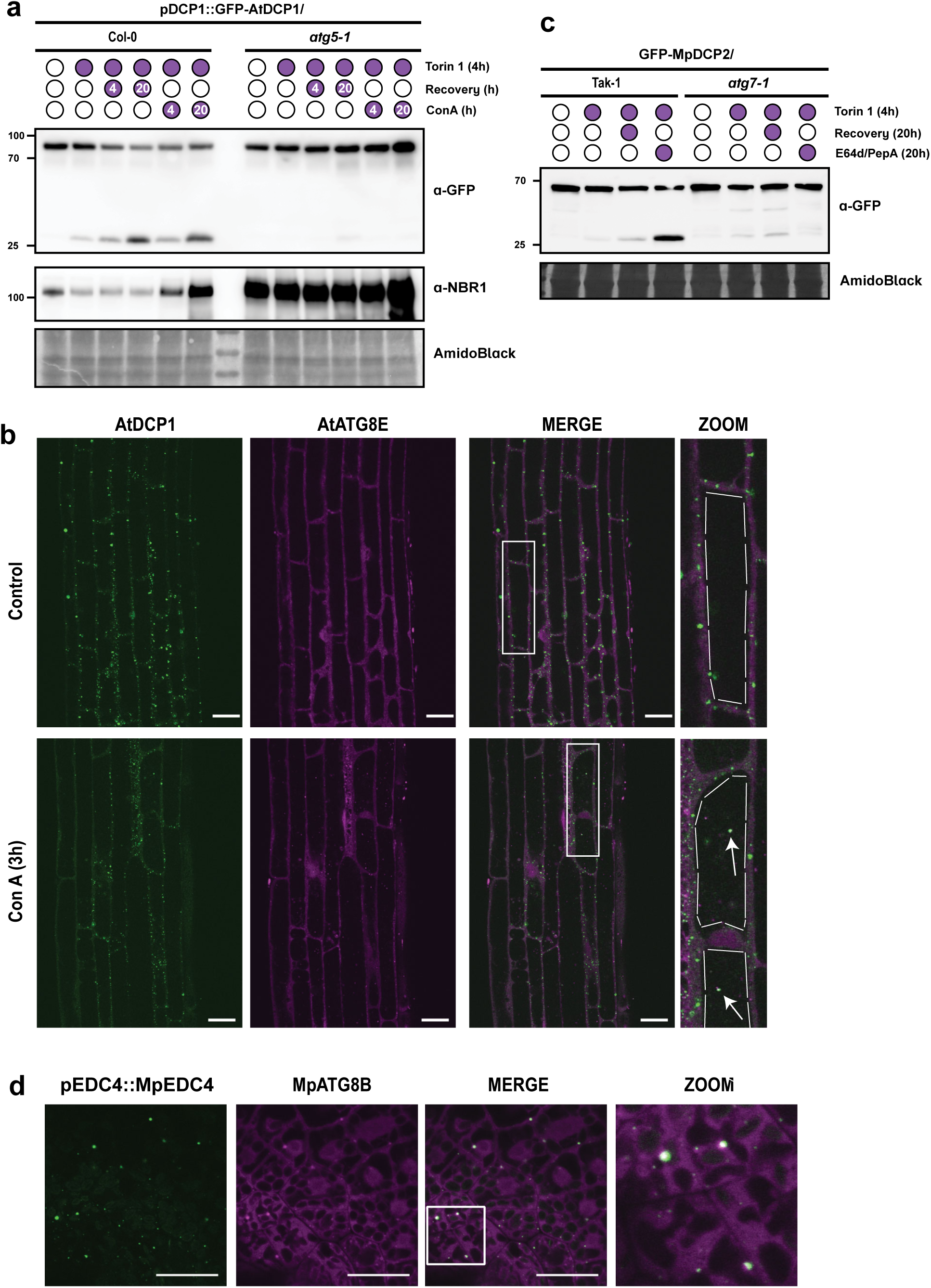
P-bodies components undergo autophagy degradation in plants. **a, AtDCP1 undergoes vacuolar degradation in an autophagy-dependent manner.** Western blot showing YFP-AtDCP1 flux assay and endogenous NBR1 level in Col-0 or *atg5-1* mutant. 7-days old *A. thaliana* seedlings were incubated in either DMSO or 9 μM Torin-1-containing media (1/2 MS, MES, 1% sucrose) for 4 h, after which plants were either harvested or recovered for additional 4 h or 20 h with either fresh media (Rec) or media containing 1 μM concanamycin A (ConA) before harvesting. Total protein levels were analyzed by staining with AmidoBlack. A second independent biological replicate is shown in Fig. S2B. **b, AtDCP1 co-localizes with AtATG8-decorated autophagosomes upon concanamycin A treatment**. Confocal microscopy images of *A. thaliana* seedlings co-expressing YFP-AtDCP1 with mCherry-ATG8E. Seedlings were incubated in 1/2 MS, MES, 1% sucrose media. Representative images of minimum 10 biological replicates are shown. Zoomed panels show enlarged sections of the white-boxed areas highlighted in the merge panel. Scale bar, 20 μm. **c, MpDCP2 undergoes vacuolar degradation in an autophagy-dependent manner.** Western blot showing GFP-MpDCP2 flux assay in Tak-1 or *atg7-1* mutant. 6-days old *M. polymorpha* gemmae were incubated in either DMSO or 12 μM Torin-1-containing media (0.5 Gamborg, MES, 1% sucrose) for 4 h, after which plants were either harvested or recovered for additional 20 h with either fresh media (Rec) or media containing 10 μM E64d and 10 μM pepstatin A (E64d/PepA) before harvesting. Total protein levels were analyzed by staining with AmidoBlack. Two independent biological replicates are shown in Fig. S2C-D **d, MpEDC4 co-localizes with MpATG8-decorated autophagosomes.** Confocal microscopy images of 2-days old *M. polymorpha* thallus cells co-expressing mScarlet-MpATG8B with GFP-MpEDC4 under minimal EDC4 promoter. Thalli were incubated in 0.5 Gamborg, MES, 1% sucrose media. Representative images of minimum 10 biological replicates are shown. Zoomed panels show enlarged sections of the white-boxed areas highlighted in the merge panel. Scale bar, 20 μm.

**Figure S2.**
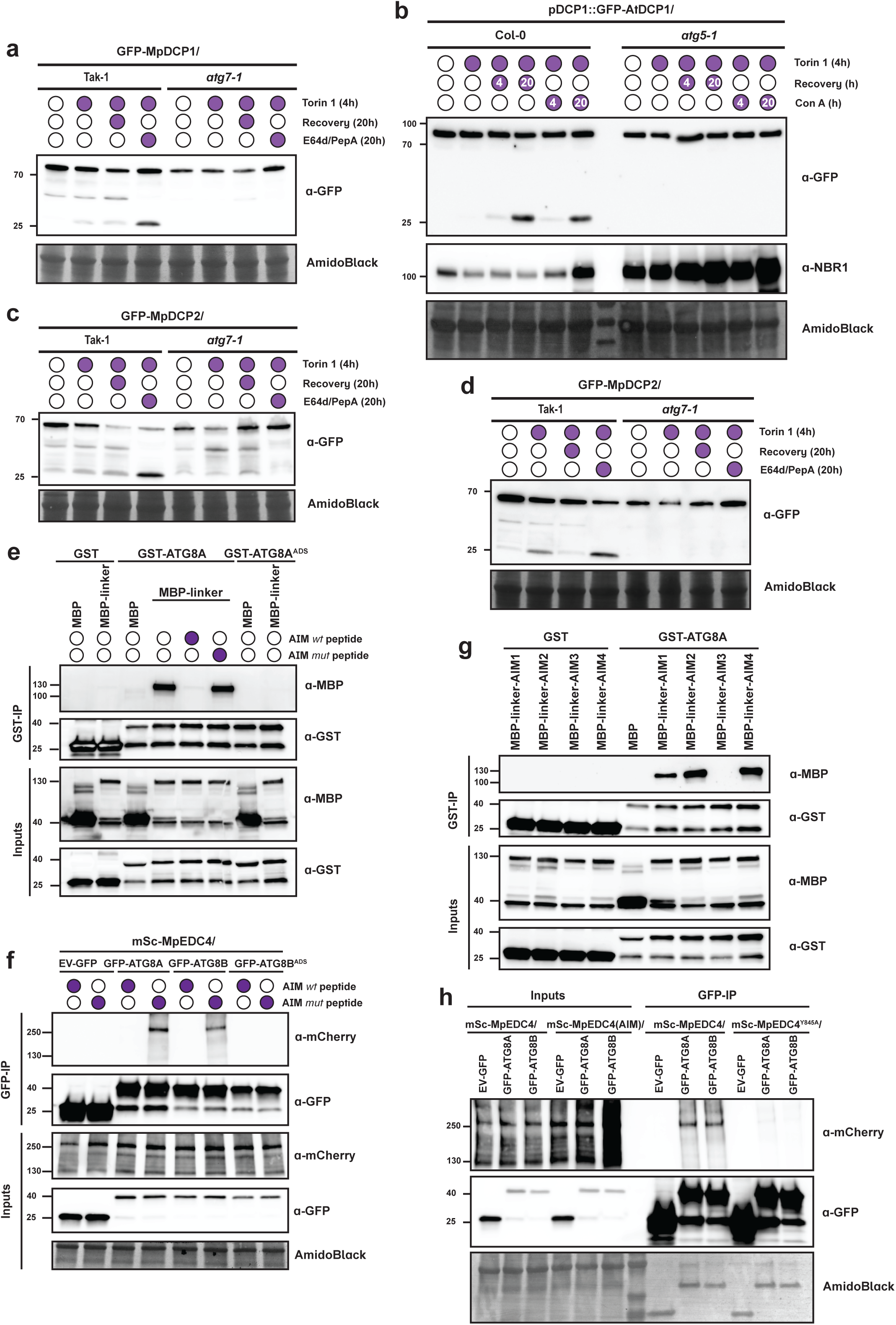
Biological replicates for Fig. 1 and Fig. S1. **a,** Second biological replicate for Fig. 1A. **b,** Second biological replicate for Fig. S1A. **c-d,** Two biological replicates for Fig. S1C. **e,** Second biological replicate for Fig. 1E. **f,** Second biological replicate for Fig. 1F. **g,** Second biological replicate for Fig. 1H. **h,** Second biological replicate for Fig. 1I

**Figure S3.**
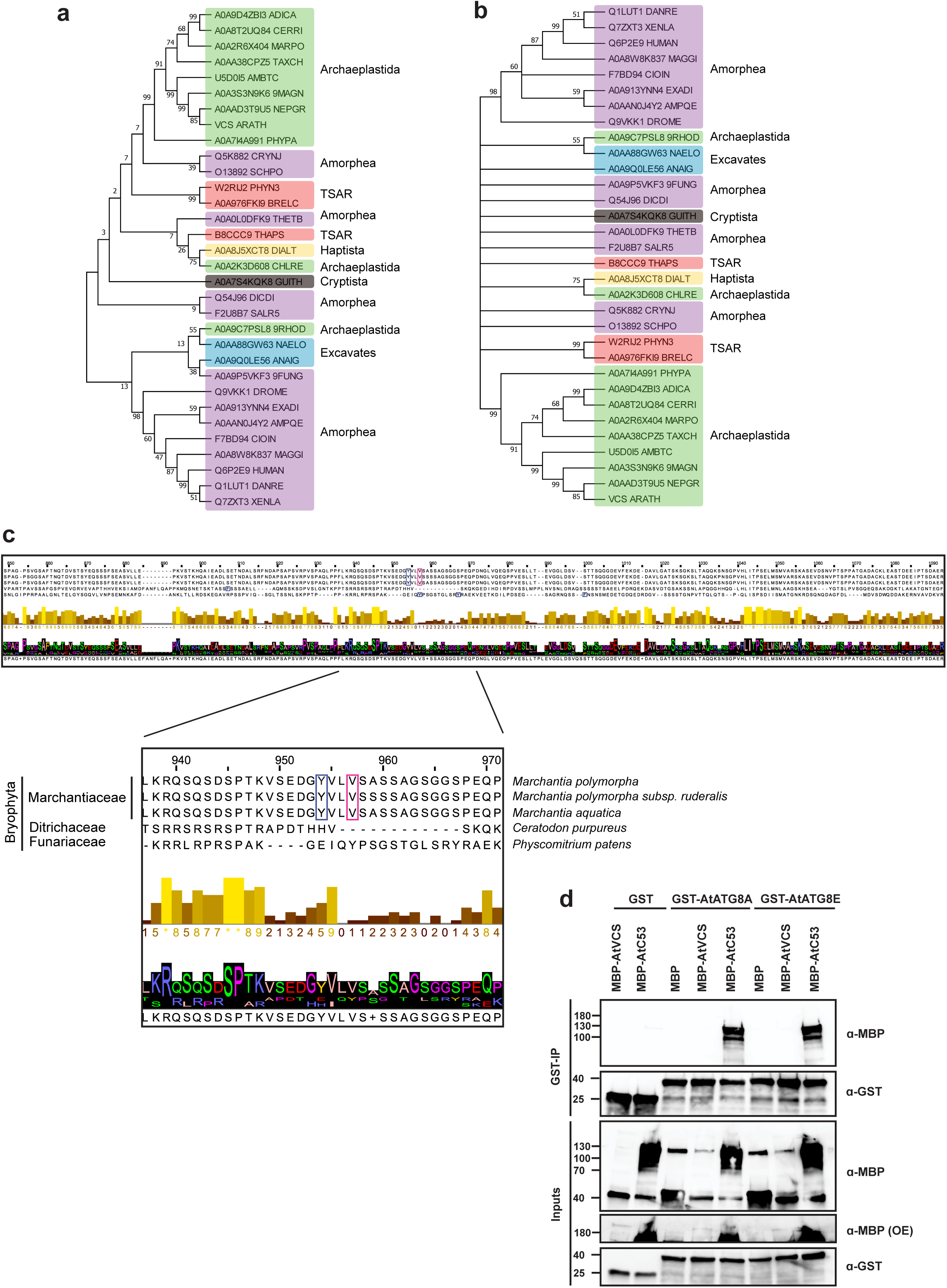
Phylogenetic analysis of EDC4. **a, Maximum likelihood phylogenetic tree of EDC4 from 32 different species across Eukaryotes**. **b, Maximum likelihood phylogenetic tree with bootstep support (n = 500 replicates) of EDC4 from 32 different species across Eukaryotes**. **c, Multiple sequence alignment of EDC4 in Bryophyta revealed that only Marchantiaceae EDC4s have canonical AIM motif withing the disordered region**. d, AtVCS does not interact with AtATG8A or AtATG8E. *In vitro* pulldown assay showing that *Arabidopsis thaliana* ortholog of MpEDC4, AtVCS, does not interact with AtATG8A or AtATG8E. Bacterial lysates containing recombinant protein were mixed and pulled down with GST magnetic agarose beads. Input and bound proteins were immunoblotted with anti-GST and anti-MBP antibodies. A second biological replicate is shown in Fig. S6F.

**Figure S4.**
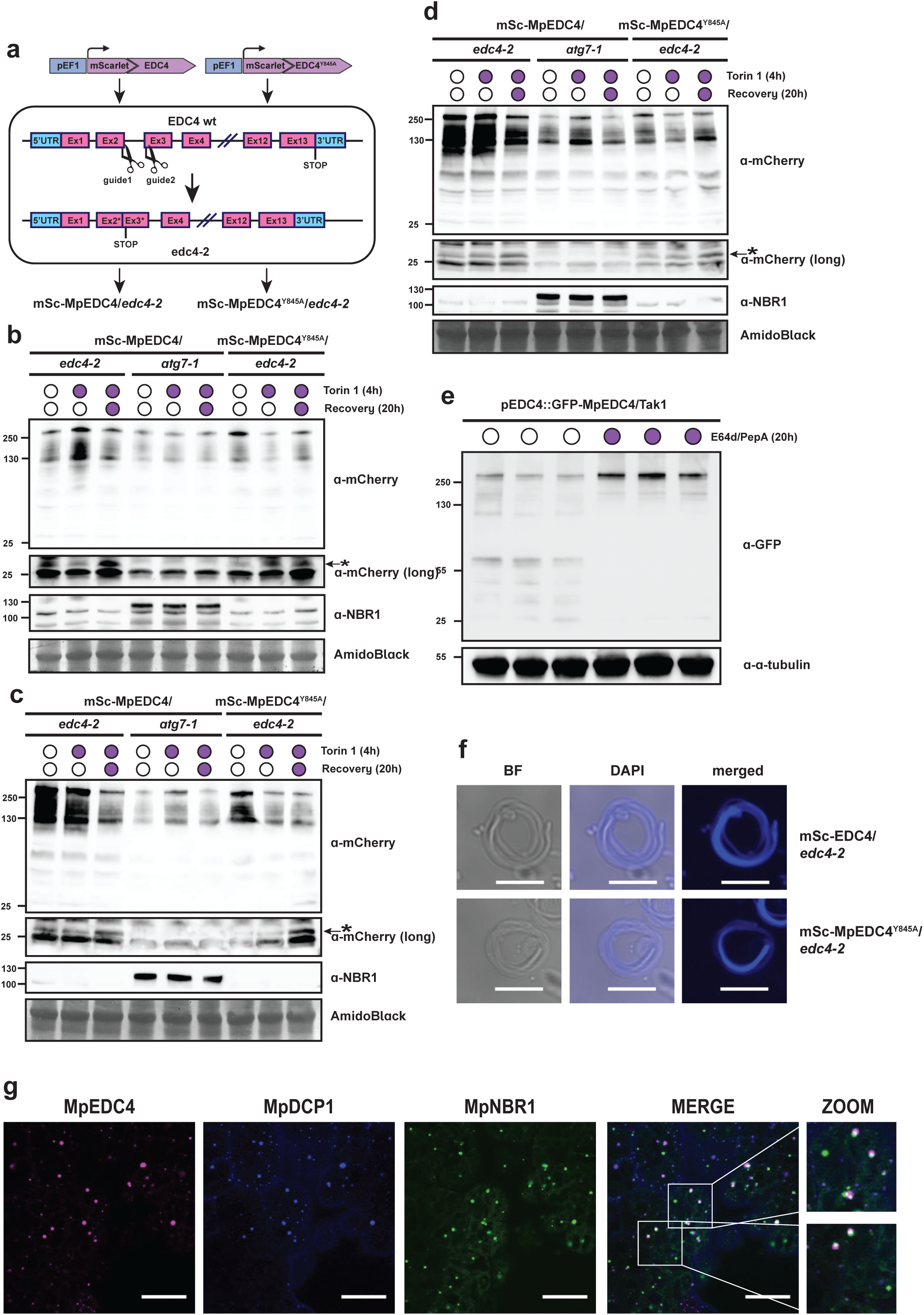
EDC4-independent autophagy of P-bodies. **a, *edc4* CRISPR-Cas9 mutant generation.** *M. polymorpha* male plants (Tak-1) expressing mSc-MpEDC4 *wt* and mSc-MpEDC4^Y845A^ were crossed to female plants (Tak-2). Obtained spores were collected and used to transformation with guide RNAs targeting intron-exon junctions in order to create premature stop codon without affecting coding sequence of mScarlet-EDC4 construct. **b-d,** Three biological replicates for Fig. 3B. **e, GFP-tagged MpEDC4 under minimal promoter is stabilized upon treatment with vacuolar protease inhibitors E64d/pepstatin A**. Western blot showing GFP-MpEDC4 in 6-days old *M. polymorpha* gemmae. Gemmae were incubated 20 h in either DMSO or media containing 10 μM E64d and 10 μM pepstatin A (E64d/PepA) before harvesting. Total protein levels were analyzed by staining with AmidoBlack. Three independent biological replicates are shown together. **f, Plants expressing both mScarlet-tagged MpEDC4 *wt and* MpEDC4^Y845A^ in *edc4-2* mutant background produced intact spermatozoids.** Mature antheridiophores were transferred to the petri dish and supplemented with 500 uL of sterile water to release spermatozoids. Spermatozoids were stained with 4′,6-diamidino-2-phenylindole (DAPI). **g, P-body components, MpEDC4 and MpDCP1, partially co-localize with MpNBR1.** Confocal microscopy images of 2-days old *M. polymorpha* thallus cells co-expressing GFP-MpNBR1 with mScarlet-MpEDC4 and MpDCP1-mKalama-3xflag. Thalli were incubated in 0.5 Gamborg, MES, 1% sucrose media. Representative images of minimum 10 biological replicates are shown. Zoomed panels show enlarged sections of the white-boxed areas highlighted in the merge panel. Scale bar, 20 μm.

**Figure S5.**
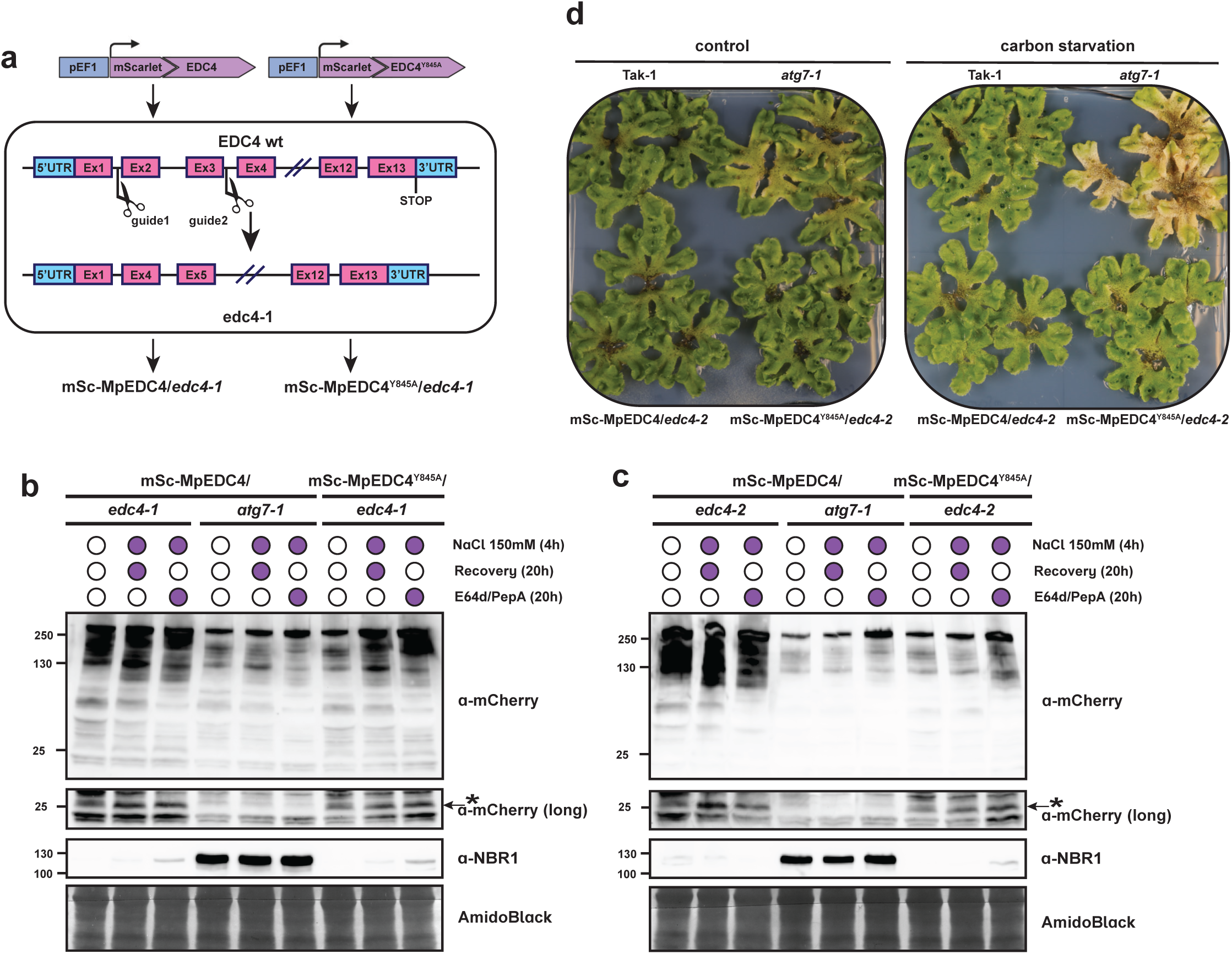
EDC4-independent autophagy of P-bodies. **a, Alternative *edc4* CRISPR-Cas9 mutant generation.** *M. polymorpha* male plants (Tak-1) expressing mScarlet-tagged MpEDC4 *wt* and MpEDC4^Y845A^ were crossed to female plants (Tak-2). Obtained spores were collected and used to transformation with guide RNAs targeting intron-exon junctions in order to cut exon2 and exon3 of *Edc4* gene without affecting coding sequence of mScarlet-EDC4 construct. **b-c, MpEDC4^Y845A^ undergoes vacuolar degradation in *edc4-/-* plants (*edc4-1* and *edc4-2*) upon high salt treatment.** Western blot showing mScarlet-MpEDC4 *wt* and MpEDC4^Y845A^ flux assay and endogenous NBR1 level in *edc4-1* **(b)** or *edc4-2* **(c)** mutant backgrounds. As control, plants expressing mScarlet-tagged MpEDC4 *wt* in *atg7-1* mutant were used. 6-days old *M. polymorpha* gemmae were incubated in either fresh media or 150 mM sodium-chloride (NaCl) containing media (0.5 Gamborg, MES, 1% sucrose) for 4 h, after which plants were either harvested or recovered for additional 20 h with either fresh media (Rec) or media containing 10 μM E64d and 10 μM pepstatin A (E64d/PepA) before harvesting. Total protein levels were analyzed by staining with AmidoBlack. *: free mScarlet band. **d,** Second biological replicate for Fig. 3D.

**Figure S6.**
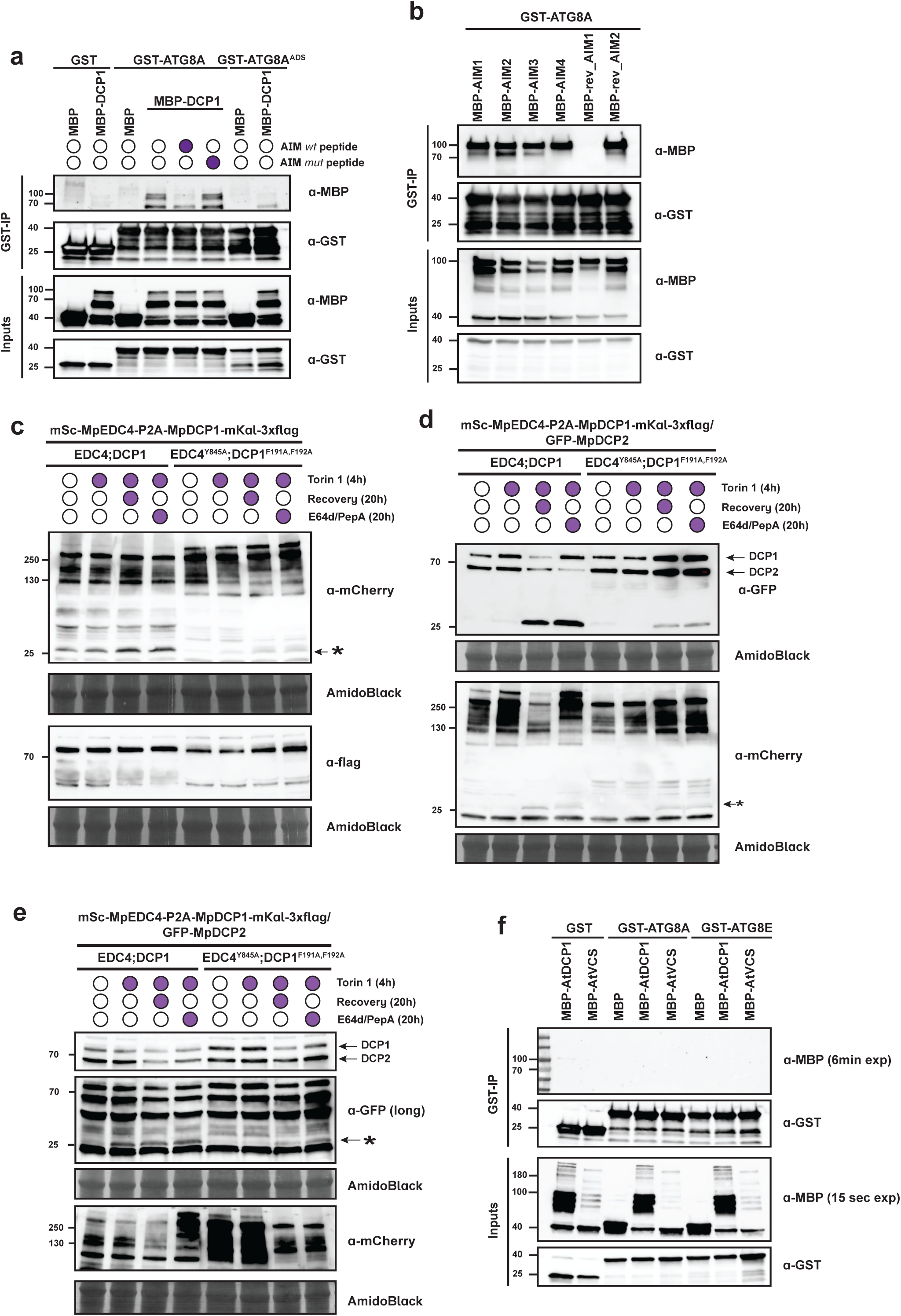
MpDCP1 interacts with ATG8. **a,** Second biological replicate for Fig. 4B. **b,** Second biological replicate for Fig. 4E. **c,** Second biological replicate for Fig. 4H. **d-e,** Two biological replicates for Fig. 4I. **f,** Second biological replicate for Fig. S3D and S7D.

**Figure S7.**
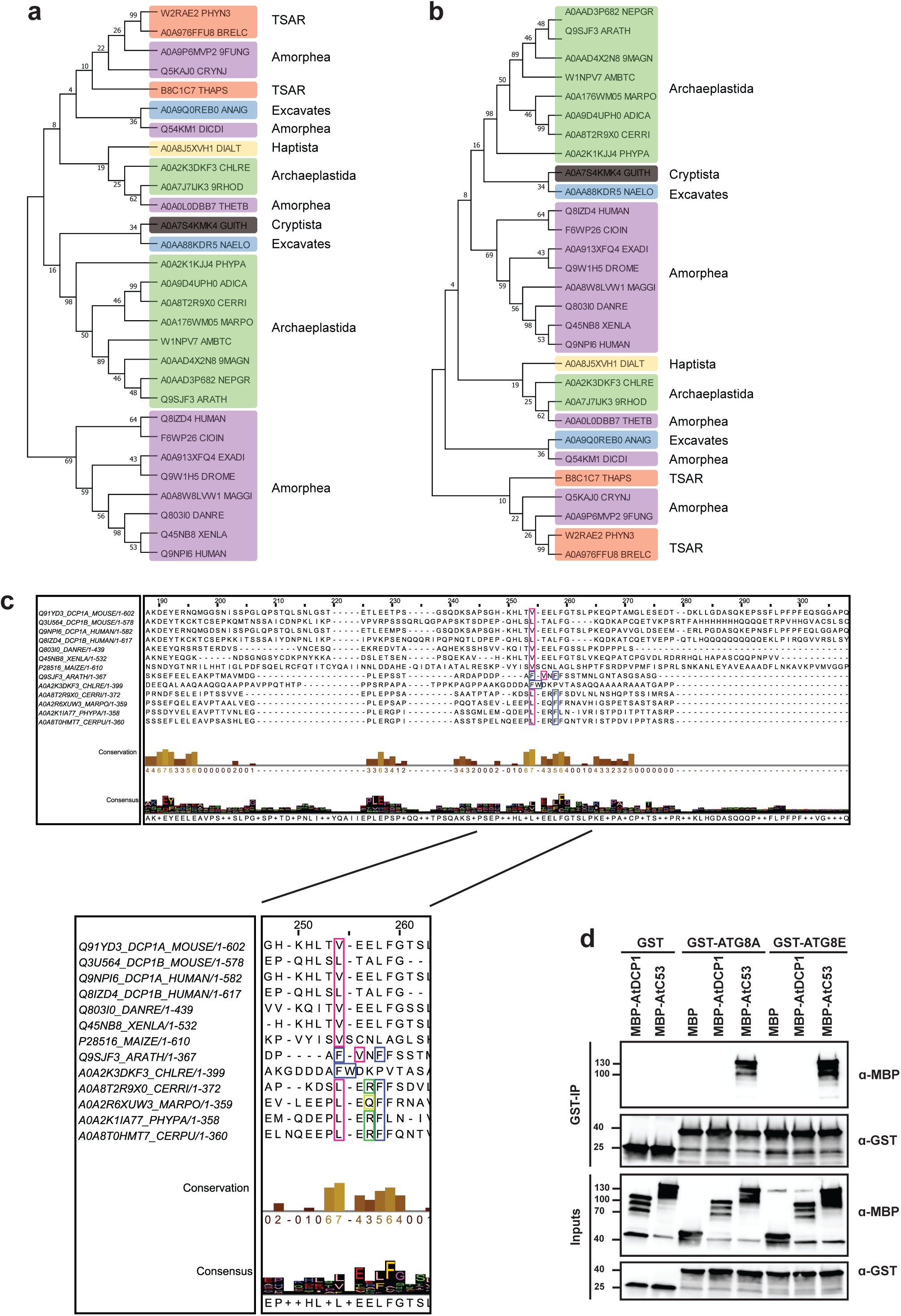
Phylogenetic analysis of DCP1. **a-b, Maximum likelihood phylogenetic tree of DCP1 from 29 different species across Eukaryotes**. **b, Maximum likelihood phylogenetic tree with bootstep support (n = 500 replicates) of DCP1 from 29 different species across Eukaryotes**. **c, Multiple sequence alignment of DCP1 in various species around reverse AIM motif**. d, AtDCP1 does not associate with AtATG8A or AtATG8E *in vitro*. *In vitro* pulldown assay showing *Arabidopsis thaliana* ortholog of MpDCP1, AtDCP1, does not interact with AtATG8A or AtATG8E isoforms. Bacterial lysates containing recombinant protein were mixed and pulled down with GST magnetic agarose beads. Input and bound proteins were immunoblotted with anti-GST and anti-MBP antibodies. A second biological replicate is shown in Fig. S6E

**Figure S8.**
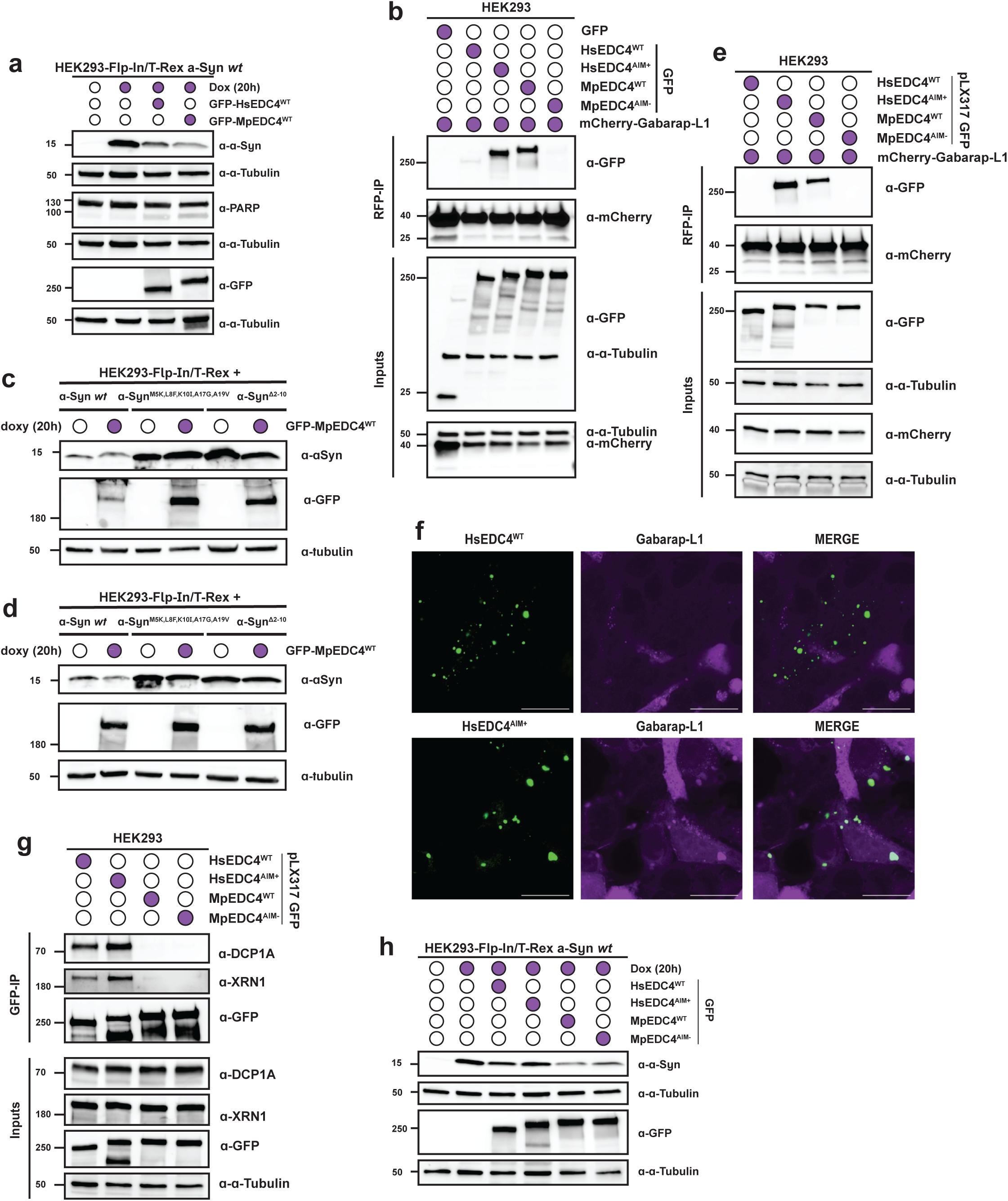
MpEDC4 interacts with human ATG8 ortholog and can be delivered to the lysosomes. **a,** Second biological replicate for Fig. 5B. **b,** Second biological replicate for Fig. 5C. **c-d,** Two biological replicates for Fig. 5D. **e, Chimeric HsEDC4 HsEDC4^AIM+^ associates with HsATG8 ortholog, Gabarap-L1 in lines stably expressing GFP versions of EDC4.** *In vivo* RFP-Trap Co-IP in HEK293T cells showing that both GFP-tagged HsEDC4^AIM+^ chimeric protein and MpEDC4^WT^ associates with mCherry-tagged Gabarap-L1, unlike HsEDC4^WT^ or MpEDC4^AIM-^. Protein extracts were immunoblotted with anti-GFP and anti-mCherry antibodies. Anti-α-tubulin antibodies were used as loading control for inputs. **f, GFP-tagged HsEDC4^AIM+^ co-localizes with mCherry-Gabarap-L1.** Confocal microscopy images of HEK293T cells co-expressing various GFP-EDC4 with mCherry-Gabarap-L1. Representative images of minimum 10 biological replicates are shown. Zoomed panels show enlarged sections of the white-boxed areas highlighted in the merge panel. Scale bar, 20 μm. **g,** Second biological replicate for Fig. 5G. **h,** Second biological replicate for Fig. 5H

## References

1. Parker, R. & Sheth, U. P Bodies and the Control of mRNA Translation and Degradation. Mol. Cell 25, 635–646 (2007).

2. Unterholzner, L. & Izaurralde, E. SMG7 Acts as a Molecular Link between mRNA Surveillance and mRNA Decay. Mol. Cell 16, 587–596 (2004).

3. Hubstenberger, A. et al. P-Body Purification Reveals the Condensation of Repressed mRNA Regulons. Mol. Cell 68, 144–157.e5 (2017).

4. Decker, C. J., Teixeira, D. & Parker, R. Edc3p and a glutamine/asparagine-rich domain of Lsm4p function in processing body assembly in Saccharomyces cerevisiae. J. Cell Biol. 179, 437–449 (2007).

5. Molliex, A. et al. Phase separation by low complexity domains promotes stress granule assembly and drives pathological fibrillization. Cell 163, 123–133 (2015).

6. Cui, Q., Liu, Z. & Bai, G. Friend or foe: The role of stress granule in neurodegenerative disease. Neuron 112, 2464–2485 (2024).

7. Buchan, J. R., Kolaitis, R. M., Taylor, J. P. & Parker, R. Eukaryotic Stress Granules Are Cleared by Autophagy and Cdc48/VCP Function. Cell 153, 1461–1474 (2013).

8. Barrow, E. R. et al. Discovery of SQSTM1/p62-dependent P-bodies that regulate the NLRP3 inflammasome. Cell Rep. 43, (2024).

9. Pan, C., Knutson, S. D., Huth, S. W. & MacMillan, D. W. C. µMap proximity labeling in living cells reveals stress granule disassembly mechanisms. Nat. Chem. Biol. 2024 1– 11 (2024) doi:10.1038/s41589-024-01721-2.

10. Gatica, D., Lahiri, V. & Klionsky, D. J. Cargo recognition and degradation by selective autophagy. Nat. Cell Biol. 20, 233–242 (2018).

11. Zaffagnini, G. & Martens, S. Mechanisms of Selective Autophagy. J. Mol. Biol. 428, 1714–1724 (2016).

12. Abdrakhmanov, A., Gogvadze, V. & Zhivotovsky, B. To Eat or to Die: Deciphering Selective Forms of Autophagy. Trends Biochem. Sci. 45, 347–364 (2020).

13. Nishimura, T. & Tooze, S. A. Emerging roles of ATG proteins and membrane lipids in autophagosome formation. Cell Discov. 2020 61 6, 1–18 (2020).

14. Stolz, A., Ernst, A. & Dikic, I. Cargo recognition and trafficking in selective autophagy. Nat. Cell Biol. 16, 495–501 (2014).

15. Birgisdottir, Å. B., Lamark, T. & Johansen, T. The LIR motif - crucial for selective autophagy. J. Cell Sci. 126, 3237–3247 (2013).

16. Rogov, V. V., et al. Atg8 family proteins, LIR/AIM motifs and other interaction modes. Autophagy Reports 2, (2023).

17. Papadopoulos, C., Kravic, B. & Meyer, H. Repair or Lysophagy: Dealing with Damaged Lysosomes. J. Mol. Biol. (2019) doi:10.1016/j.jmb.2019.08.010.

18. Lazarou, M. et al. The ubiquitin kinase PINK1 recruits autophagy receptors to induce mitophagy HHS Public Access. Nature 524, 309–314 (2015).

19. Forrester, A. et al. A selective ER-phagy exerts procollagen quality control via a Calnexin-FAM134B complex. EMBO J. 38, e99847 (2019).

20. Princely Abudu, Y., et al. NIPSNAP1 and NIPSNAP2 Act as “Eat Me” Signals for Mitophagy. Dev. Cell (2019) doi:10.1016/j.devcel.2019.03.013.

21. Trapannone, R., Romanov, J. & Martens, S. p62 and NBR1 functions are dispensable for aggrephagy in mouse ESCs and ESC-derived neurons. Life Sci. Alliance 6, (2023).

22. Lu, K., Psakhye, I. & Jentsch, S. Autophagic clearance of PolyQ proteins mediated by ubiquitin-Atg8 adaptors of the conserved CUET protein family. Cell 158, 549–563 (2014).

23. Thurston, T. L. M., Ryzhakov, G., Bloor, S., von Muhlinen, N. & Randow, F. The TBK1 adaptor and autophagy receptor NDP52 restricts the proliferation of ubiquitin-coated bacteria. Nat. Immunol. 10, 1215–1221 (2009).

24. Ravenhill, B. J. et al. The Cargo Receptor NDP52 Initiates Selective Autophagy by Recruiting the ULK Complex to Cytosol-Invading Bacteria. Mol. Cell 74, 320–329.e6 (2019).

25. Ma, X. et al. CCT2 is an aggrephagy receptor for clearance of solid protein aggregates. Cell 185, 1325–1345.e22 (2022).

26. Kraft, C., Deplazes, A., Sohrmann, M. & Peter, M. Mature ribosomes are selectively degraded upon starvation by an autophagy pathway requiring the Ubp3p/Bre5p ubiquitin protease. Nat. Cell Biol. 10, 602–610 (2008).

27. Wyant, G. A. et al. NUFIP1 is a ribosome receptor for starvation-induced ribophagy. Science *(80-.).* **360**, 751–758 (2018).

28. Parker, R. & Song, H. The enzymes and control of eukaryotic mRNA turnover. Nat. Struct. Mol. Biol. 11, 121–127 (2004).

29. Tanida, I., Minematsu-Ikeguchi, N., Ueno, T. & Kominami, E. Lysosomal turnover, but not a cellular level, of endogenous LC3 is a marker for autophagy. Autophagy 1, 84– 91 (2005).

30. Stephani, M. et al. A cross-kingdom conserved er-phagy receptor maintains endoplasmic reticulum homeostasis during stress. Elife 9, 1–105 (2020).

31. Hernández, V. S. de M. et al. Cross-species interactome analysis uncovers a conserved selective autophagy mechanism for protein quality control in plants. *bioRxiv* 2024.09.08.611708 (2024) doi:10.1101/2024.09.08.611708.

32. Noda, N. N., Ohsumi, Y. & Inagaki, F. Atg8-family interacting motif crucial for selective autophagy. FEBS Lett. 584, 1379–1385 (2010).

33. Hurley, J. H. & Schulman, B. A. Atomistic Autophagy: The Structures of Cellular Self- Digestion. Cell 157, 300 (2014).

34. Rozenknop, A. et al. Characterization of the Interaction of GABARAPL-1 with the LIR Motif of NBR1. J. Mol. Biol. 410, 477–487 (2011).

35. Chang, C. Te, Bercovich, N., Loh, B., Jonas, S. & Izaurralde, E. The activation of the decapping enzyme DCP2 by DCP1 occurs on the EDC4 scaffold and involves a conserved loop in DCP1. Nucleic Acids Res. 42, 5217–5233 (2014).

36. Suzuki, H. & Noda, N. N. Biophysical characterization of Atg11, a scaffold protein essential for selective autophagy in yeast. FEBS Open Bio 8, 110 (2017).

37. Zientara-Rytter, K. & Subramani, S. Mechanistic insights into the role of Atg11 in selective autophagy. J. Mol. Biol. 432, 104 (2019).

38. Zheng, D., Chen, C. Y. A. & Shyu, A. Bin. Unraveling regulation and new components of human P-bodies through a protein interaction framework and experimental validation. RNA 17, 1619 (2011).

39. Xie, Z. et al. Proteasome resides in and dismantles plant heat stress granules constitutively. Mol. Cell 84, 3320–3335.e7 (2024).

40. Okumura, M., Katsuyama, A. M., Shibata, H. & Maki, M. VPS37 isoforms differentially modulate the ternary complex formation of ALIX, ALG-2, and ESCRT-I. Biosci. Biotechnol. Biochem. 77, 1715–1721 (2013).

41. Xu, J., Yang, J. Y., Niu, Q. W. & Chua, N. H. Arabidopsis DCP2, DCP1, and VARICOSE form a decapping complex required for postembryonic development. Plant Cell 18, 3386–3398 (2006).

42. Norizuki, T., Minamino, N., Sato, M. & Ueda, T. Autophagy regulates plastid reorganization during spermatogenesis in the liverwort Marchantia polymorpha. Front. Plant Sci. 14, 1101983 (2023).

43. Norizuki, T., Minamino, N., Sato, M., Tsukaya, H. & Ueda, T. Dynamic rearrangement and autophagic degradation of mitochondria during spermiogenesis in the liverwort Marchantia polymorpha. Cell Rep. 39, 110975 (2022).

44. Hallacli, E. et al. The Parkinson’s disease protein alpha-synuclein is a modulator of processing bodies and mRNA stability. Cell 185, 2035–2056.e33 (2022).

45. Maldonado-Bonilla, L. D. Composition and function of P bodies in Arabidopsis thaliana. Frontiers in Plant Science vol. 5 at 10.3389/fpls.2014.00201 (2014).

46. An, H. et al. TEX264 Is an Endoplasmic Reticulum-Resident ATG8-Interacting Protein Critical for ER Remodeling during Nutrient Stress. Mol. Cell 74, 891–908.e10 (2019).

47. Nguyen, T. N. et al. Unconventional initiation of PINK1/Parkin mitophagy by Optineurin. Mol. Cell 83, 1693–1709.e9 (2023).

48. Kirkin, V. & Rogov, V. V. A Diversity of Selective Autophagy Receptors Determines the Specificity of the Autophagy Pathway. Mol. Cell 76, 268–285 (2019).

49. jetOPTIMUS® - DNA transfection reagent. https://www.polyplus-sartorius.com/products/jetoptimus.

50. Sauret-Güeto, S. et al. Systematic Tools for Reprogramming Plant Gene Expression in a Simple Model, Marchantia polymorpha. ACS Synth. Biol. 9, 864–882 (2020).

51. Lampropoulos, A. et al. GreenGate - A novel, versatile, and efficient cloning system for plant transgenesis. PLoS One 8, (2013).

52. Tsuboyama, S., Nonaka, S., Ezura, H. & Kodama, Y. Improved G-AgarTrap: A highly efficient transformation method for intact gemmalings of the liverwort Marchantia polymorpha. Sci. Reports 2018 81 8, 1–10 (2018).

53. Clough, S. J. & Bent, A. F. Floral dip: a simplified method for Agrobacterium-mediated transformation of Arabidopsis thaliana. Plant J. 16, 735–743 (1998).

54. Madeira, F. et al. The EMBL-EBI search and sequence analysis tools APIs in 2019. Nucleic Acids Res. 47, W636–W641 (2019).

55. Kumar, S., Nei, M., Dudley, J. & Tamura, K. MEGA: A biologist-centric software for evolutionary analysis of DNA and protein sequences. Brief. Bioinform. 9, 299–306 (2008).

56. Stephani, M., Picchianti, L. & Dagdas, Y. C53 is a cross-kingdom conserved reticulophagy receptor that bridges the gap betweenselective autophagy and ribosome stalling at the endoplasmic reticulum. Autophagy 17, 586–587 (2021).

57. Norizuki, T., Kanazawa, T., Minamino, N., Tsukaya, H. & Ueda, T. Marchantia polymorpha, a new model plant for autophagy studies. Front. Plant Sci. 10, 467892 (2019).

58. Thompson, A. R., Doelling, J. H., Suttangkakul, A. & Vierstra, R. D. Autophagic Nutrient Recycling in Arabidopsis Directed by the ATG8 and ATG12 Conjugation Pathways. Plant Physiol. 138, 2097–2110 (2005).

59. Sessions, A. et al. A High-Throughput Arabidopsis Reverse Genetics System. Plant Cell 14, 2985–2994 (2002).

60. Billey, E. et al. LARP6C orchestrates posttranscriptional reprogramming of gene expression during hydration to promote pollen tube guidance. Plant Cell 33, 2637– 2661 (2021).

